# scDNAm-GPT Captures Genome-wide CpG Dependencies in Single-cell DNA methylomes to Revolutionize Epigenetic Analysis

**DOI:** 10.1101/2025.02.19.638959

**Authors:** Chaoqi Liang, Peng Ye, Hongliang Yan, Peng Zheng, Jianle Sun, Yanni Wang, Yu Li, Yuchen Ren, Yuanpei Jiang, Ran Wei, Junjia Xiang, Sizhe Zhang, Linle Jiang, Weiqiang Bai, Xinzhu Ma, Tao Chen, Wangmeng Zuo, Lei Bai, Wanli Ouyang, Jia Li

## Abstract

Single-cell DNA methylomes are challenging to interpret because of sparse CpG coverage and the complexity of genome-wide sequences. We present scDNAm-GPT, a universal foundation model that uses context-aware CpG tokenization, a Mamba backbone, and cross-attention to capture both local and global DNA methylation patterns. Trained on over one million single cells from 35 human and mouse tissues, scDNAm-GPT enables accurate cell clustering, zero-shot prediction of CpG effects on gene expression, improved trajectory inference, and reference-free deconvolution of cell types from cell-free DNA. The model hierarchically learns regulatory features, and its attention maps highlight functionally relevant regions, demonstrating high biological interpretability. These results establish scDNAm-GPT as a scalable and generalizable framework for single-cell epigenomic analysis, offering new opportunities to dissect epigenetic regulation in health and disease. Code is available at GitHub (https://github.com/ChaoqiLiang/scDNAm-GPT).

## Introduction

Beyond the four canonical DNA bases (A, T, C, and G), DNA methylation is regarded as the “fifth base” and represents one of the most prevalent epigenetic modifications in mammals. It acts in concert with other epigenetic layers, such as histone modifications and chromatin architecture, to regulate gene expression during development, maintain cellular identity, and mediate responses to environmental stimuli. Aberrant DNA methylation disrupts epigenomic integrity, driving transcriptional dysregulation and contributing to diverse diseases, particularly cancer. Besides, tumor cell-derived cell-free DNA fragments carrying distinctive DNA methylation signatures circulate in body fluids, making DNA methylation as a powerful biomarker for disease detection and monitoring.

The advent of single-cell whole-genome bisulfite sequencing (scWGBS) (*1, 2*) has provided an unprecedented opportunity to map genome-wide DNA methylation at single-cell and single-nucleotide resolution, enabling precise characterization of cell-to-cell variability, cellular trajectories, and disease-associated DNA methylation variations. However, realizing this potential is hindered by several analytical limitations. Prevailing analytical approaches rely on DNA methylation features in predefined genomic bins (e.g., 100-kb or 5-kb windows), obscuring fine-scale patterns and long-range CpG dependencies and limiting biological interpretability. Differential methylation analysis across conditions further relies on customized, non-standardized methods, complicating cross-study harmonization. Recent bulk DNA methylation models (*3*) require manual feature selection, introducing bias and restricting their generalizability across tasks.

These widespread analytical limitations necessitate a new computational paradigm. Inspired by the transformative success of foundation models in single-cell transcriptomics and proteomics, we aimed to extend this approach to DNA methylation, which, unlike downstream readouts of gene regulation, represents a fundamental upstream layer of epigenetic control. Although scWGBS captures this crucial upstream information, no foundation model exists for single-cell DNA methylome analysis to date. This gap stems from two primary hurdles. First, while transformer-based generative models such as scBERT (*4*), scGPT (*5*), and scFoundation (*6*) have demonstrated remarkable success in other domains, their quadratic self-attention cost prevents direct application to the ultra-long sequences inherent in single-cell DNA methylome data. Second, scWGBS data present unique computational challenges—with over 20 million CpG sites per genome, extreme sparsity, and high stochasticity—that overwhelm conventional machine learning approaches. Addressing these hurdles requires a new framework capable of efficiently handling long-sequence inputs, automatically learning genome-wide CpG context, and dynamically prioritizing key CpG sites from highly sparse data to answer diverse biological questions.

Here, we developed scDNAm-GPT, the first foundation model for genome-wide single-cell DNA methylomes (up to 100 million base pairs), distinguished from existing single-cell models by its Mamba architecture with a well-designed tokenizer and prediction modules, enabling efficient analysis of ultra-long methylome sequences—far exceeding existing models such as Evo2 (one million base pairs) (*7*). It integrates three key components: i) The CpG-centered 6-mer tokenization that models CpG with surrounding DNA; ii) Mamba’s Selective State Space Models (SSMs) (*8*) that handle ultra-long sequences; and iii) The cross-attention heads that integrate nucleotide-level context, enabling scDNAm-GPT to model both local and distal dependencies. scDNAm-GPT was pretrained on one million single-cell DNA methylomes from 35 tissues across human and mouse (*9–12*), with each cell containing on average 1.3 million CpG sites and some reaching up to 20 million. This large-scale pretraining enables scDNAm-GPT to learn diverse and generalizable DNA methylation features across species and biological contexts, without needing handcrafted feature engineering.

scDNAm-GPT demonstrates exceptional performance across a wide range of downstream tasks, breaking through core challenges in single-cell epigenetics to unlock new biological insights and provide a powerful foundation for clinical applications. Specifically, scDNAm-GPT achieves new state-of-the-art performance in cell clustering, cell annotation, gene expression prediction, and pseudotime trajectory inference, while demonstrating strong generalizability to unseen datasets through zero-shot, interpretable modeling of developmental trajectories. Further, scDNAm-GPT accurately predicts the regulatory effects of individual CpG DNA methylation states on gene expression at single-cell resolution, providing a dynamic and reversible view of epigenetic regulation. Beyond fundamental discovery, scDNAm-GPT enables reference-free cfDNA deconvolution for non-invasive oncology, underscoring its translational potential for DNA methylation-based clinical applications. In general, scDNAm-GPT provides a scalable and generalizable framework for single-cell DNA methylome analysis, delivering interpretable regulatory insights and enabling translational applications from basic epigenomic discovery to cfDNA-based cancer detection, opening new avenues to decode the regulatory logic of development, disease, and cellular heterogeneity.

## Results

### scDNAm-GPT

We present scDNAm-GPT, the first CpG language model tailored for accurate and scalable interpretation of single-cell whole genome DNA methylation data, which effectively captures both local sequence context and long-range interactions among CpG sites. scDNAm-GPT leverages an innovative CpG-centered tokenization design, the Mamba framework, and cross-attention heads to process over 20 million CpG sites and more than 80 million nucleotides of surrounding CpG sites in a single cell, enabling efficient modeling and analysis of single-cell DNA methylation data.

The pretraining dataset includes 1 million single-cell DNA methylation profiles from human and mouse, covering 35 distinct cell types, including 400,000 single-cell DNA methylomes from the mouse brain and 600,000 from various human tissues. The total pretraining dataset contains 1.2 trillion CpG sites, with an average of 1.2 million CpG sites per cell, and some cells covering more than 20 million CpG sites (Fig. 1A). In designing the tokenization approach for methylation data, we considered the impact of DNA background information surrounding CpG sites on methylation (*13*). Thus, we innovatively adopted a CpG-centered k-mer design. Experimental results showed that the 6-mer representation can capture most DNA background information within the same GPU memory constraints (Fig. S1). Specifically, each CpG site and its surrounding nucleotides form a 6-mer CpG motif in the format “*XXC(M)GXX*”, where “*X*” represents any nucleotide and “*M*” indicates the methylation state. We treat each motif as a “word”, enabling the construction of a CpG language for downstream tasks. Special tokens *[BOS]* and *[SEP]* are introduced to mark dataset boundaries, enabling effective management of variable-length sequences (Fig. 1B). A key innovation is the introduction of a novel “CpG fuzzy embedding” approach, where each CpG site’s embedding is a weighted sum of methylation and unmethylation motifs, reflecting their respective methylation rates to preserve probabilistic epigenetic information (Fig. 1B). The design of fuzzy tokens enables the model to flexibly handle both single-cell and bulk methylation data, ensuring seamless compatibility in downstream tasks and enhancing the model’s broad applicability and flexibility.

**Fig. 1.**
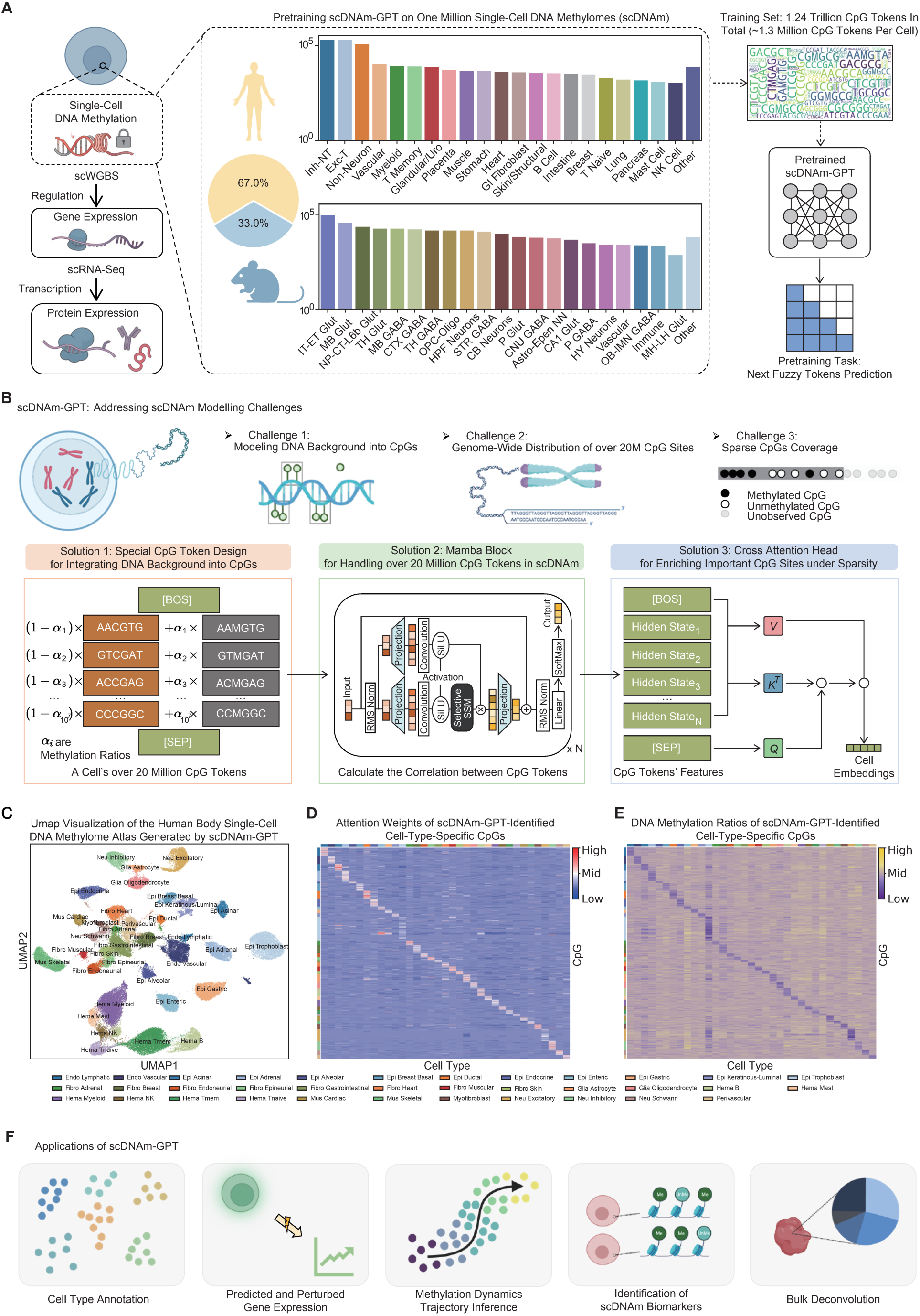
scDNAm-GPT: a foundation model for single-cell DNA methylomes and its applications. (A) We pretrain scDNAm-GPT on ∼1 million single-cell whole-genome bisulfite sequencing (scWGBS) profiles (≈1.24 trillion CpG tokens in total; ∼1.3 million CpG tokens per cell) using a next-fuzzy-token prediction objective, and use the resulting hidden state with cross attention heads to enable downstream tasks. (B) Key challenges in scDNAm modeling—i) integrating DNA background into CpGs; ii) genome-wide distribution across >20 million CpG sites; and iii) sparse coverage—are addressed by three design choices: Solution 1, a CpG-aware tokenization that embeds local DNA context; Solution 2, Mamba blocks to process ultra-long sequences; Solution 3, a cross-attention head that enriches informative CpG sites under sparsity. (C) UMAP of a human-body single-cell DNA-methylome atlas generated by scDNAm-GPT, organizing major epithelial, fibroblast, endothelial, immune, glial/neuronal, muscle, and perivascular populations across organs. (D) Attention-based identification of cell-type-specific CpG sites by scDNAm-GPT. (E) DNA methylation ratios at CpG sites highlighted by scDNAm-GPT attention weights demonstrate cell-type-specific epigenetic patterns. (F) Applications of scDNAm-GPT: (1) cell-type annotation across tissues, (2) Prediction of gene expression, (3) dynamic trajectory and pseudotime inference, (4) biological interpretability via attention-highlighted CpGs and locus-level patterns, and (5) bulk deconvolution of complex samples.

scDNAm-GPT innovatively adopts the cutting-edge Mamba architecture (*8*), specifically designed to handle the immense scale and complexity of scWGBS data, including millions of CpG words across ultra-long genomic sequences. Leveraging Selective State Space Models (SSMs) and a modular, scalable design, Mamba excels in computational efficiency through dynamic computation graphs and advanced memory optimization, minimizing memory usage while maximizing performance even on massive and sparse datasets.

The pretraining stage employs a novel “next fuzzy token prediction” strategy, an extension of traditional next token prediction tasks (*14*) tailored for DNA methylation data. At each CpG site, the model simultaneously predicts both the methylation tokens and unmethylation tokens, with the loss weights dynamically adjusted based on the methylation rate and its complement (1-methylation rate). This design effectively captures the probabilistic nature of CpG methylation, enabling the model to learn nuanced epigenetic patterns from sparse and complex single-cell data. In the finetuning stage, the model’s defining feature, Cross-Attention, allows the final hidden state of the *[SEP]* token to dynamically attend to the hidden states of all CpG sites, integrating both local and global dependencies to generate robust single-cell representations (Fig. 1B).

We next applied scDNAm-GPT to the pretraining data and generated the largest single-cell DNA methylome atlas to date, visualized in a single UMAP (Fig. 1C). The atlas revealed clear separation of major tissues and fine-grained resolution of distinct cell types, underscoring the ability of scDNAm-GPT to capture biologically meaningful DNA methylation patterns at single-CpG resolution. By leveraging the mamba and cross-attention modules of scDNAm-GPT, we identified the top 0.1% CpGs with the highest attention scores in each tissue/cell type, yielding 35 distinct tissue- and cell-type-specific DNA methylation signatures from ∼500,000 human single-cell methylomes (Fig. 1D). Notably, the DNA methylation ratios and attention scores of these CpG sites showed strong concordance across the 35 tissues and cell types (Fig. 1D, 1E).

Together, this atlas represents an unprecedented resource for dissecting cell-type-specific DNA methylation patterns and provides a foundation for functional and translational analyses of single-cell methylomes in diverse downstream tasks, such as cell clustering and annotation, trajectory inference, disease marker identification, deconvolution, and so on (Fig. 1F).

### Benchmarking the efficiency, scalability, and generalizability of scDNAm-GPT

To systematically evaluate the capabilities of scDNAm-GPT, we conducted comprehensive benchmarking across multiple biological contexts, assessing inference speed, training efficiency, predictive accuracy, and cross-tissue generalization. Five datasets were used: a human brain dataset from pretraining corpus (*11*), and four independent datasets—human developing brain cells (*15*), colorectal cancer (CRC) (*16*), and early embryonic development in both human (*17*) and mouse (*18*). These unseen datasets enabled a rigorous assessment of the model’s generalization ability. All models compared in this study, including Transformer, Mamba, and scDNAm-GPT, were pretrained on the same large-scale single-cell DNA methylation dataset (comprising 1 million cells) to ensure a fair comparison.

To benchmark the performance of our architecture on predicting cell types, the above five datasets are used. Our results demonstrate that Mamba consistently achieves more than 70% in mean accuracy, F1-score, and recall—representing an improvement of approximately 15% over Transformers in the human adult brain dataset (Fig. 2A-C). While Mamba offers an efficient backbone for modeling long-range dependencies in genomic sequences, it lacks an inherent mechanism to prioritize biologically informative regions—an essential feature for interpreting epigenetic landscapes. To overcome this limitation, scDNAm-GPT incorporates a lightweight cross-attention module on top of the Mamba architecture, enabling selective enhancement of key CpG site representations. As an illustrative example of the cross-attention mechanism, Fig. S2 presents the cross-attention landscapes of scDNAm-GPT across normal tissue and four lesion types in the CRC dataset. This visualization highlights the model’s ability to selectively prioritize variable CpG sites, enhancing their differential representation across normal and lesions. This architectural refinement further improves performance, achieving over 80% in accuracy, F1-score, and recall, representing a substantial performance gain of approximately 15% over Mamba in the five datasets. Importantly, validation on unseen datasets highlights the expanded generalizability of scDNAm-GPT, demonstrating its robustness across diverse single-cell DNA methylation profiles (Fig. 2A, 2D).

**Fig. 2.**
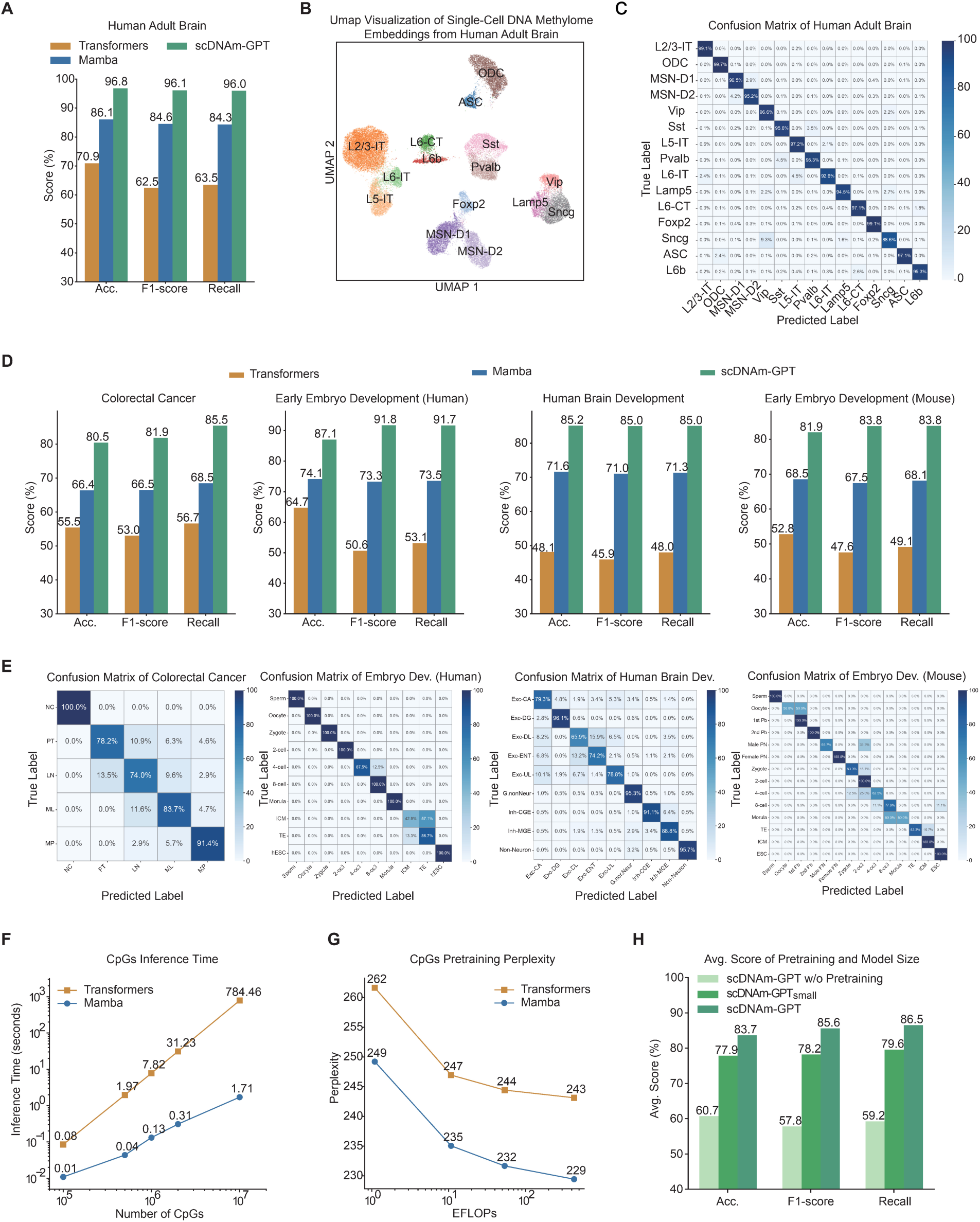
Benchmarking scDNAm-GPT for efficient, scalable, and accurate modeling of single-cell DNA methylation. (**A**) Performance comparison of Transformer, Mamba, and scDNAm-GPT on adult brain cell classification, evaluated by accuracy, F1-score, and recall. (B) UMAP of adult brain cell embeddings generated by scDNAm-GPT. (**C**) Confusion matrix on the adult brain test set, showing prediction accuracy across 15 cell types. (**D**) Performance comparison across four datasets—colorectal cancer, early embryo development (human and mouse), and human brain development—evaluated by accuracy, precision, and F1-score. scDNAm-GPT consistently outperforms both baselines. (**E**) Confusion matrices of scDNAm-GPT on the test sets of the four datasets, illustrating classification accuracy and error patterns across cell or tissue types. (**F**) Inference time of Transformer and Mamba models as a function of CpG input sequence length. Mamba scales linearly and maintains low latency even for 10⁷ CpGs, while Transformer latency increases exponentially. (**G**) Relationship between training compute (EFLOPs, 1 × 10¹⁸ floating-point operations) and model perplexity on fuzzy next-token prediction over CpG sequences. Mamba consistently achieves lower perplexity than Transformer. (**H**) Effect of model depth and pretraining on generalization. Average performance across four unseen datasets for scDNAm-GPT w/o pretraining, scDNAm-GPT_small_ (4-layer with pretraining), and scDNAm-GPT (8-layer with pretraining). Both larger model size and pretraining enhance downstream accuracy.

To assess the impact of cell number on the performance of scDNAm-GPT, we systematically evaluated classification accuracy across five datasets (Fig. 2A, 2D): the human early embryo dataset (10 developmental stages; trained on 195 cells, tested on 85), mouse early embryo (14 cell types; trained on 168 cells, tested on 72), CRC (5 lesion categories; trained on 895 cells, tested on 384), developing human brain (9 major cell types; trained on 4,096 cells, tested on 1,759), and adult human brain (15 cell types; trained on 116,643 cells, tested on 38,881 cells). The confusion matrices reveal that scDNAm-GPT achieves accuracies of 87.1% (human early embryo), 81.9% (mouse early embryo), 80.5% (CRC), and 85.2% (developing human brain), 96.8% (adult human brain) (Fig. 2A, 2C, 2D, 2E), demonstrating robust performance across datasets with training cell counts ranging from fewer than 300 to over 5,000.

We next assessed computational efficiency using the pretraining dataset. Mamba exhibited linear scalability in inference speed, processing over 10 million CpGs in under two seconds per cell, while Transformer latency increased exponentially with sequence length (Fig. 2F). Furthermore, in terms of training efficiency, Mamba consistently achieved lower perplexity than the Transformer across all compute budgets (Fig. 2G), highlighting its superior suitability for modeling ultra-long genomic sequences. To assess the impact of pretraining, we compared the performance of scDNAm-GPT with that of a randomly initialized counterpart trained from scratch under identical conditions. The pretrained model demonstrated a substantial improvement in performance, yielding gains of 23.0% in accuracy, 27.8% in F1-score, and 27.3% in recall, underscoring the advantages of large-scale DNA methylation pretraining. We further evaluated the impact of model depth on performance. Compared to the 4-layer configuration (scDNAm-GPT_small_), the 8-layer scDNAm-GPT model yields an additional 5.4% to 7.8% improvement in accuracy, F1-score, and recall on the four unseen datasets (Fig. 2H). These findings underscore the advantages of deeper architectures in capturing complex DNA methylation patterns.

scDNAm-GPT integrates Mamba and Cross-Attention to efficiently model single-cell DNA methylation data. Pretraining and the deeper architecture improve accuracy and generalizability remarkably, while Mamba ensures fast, scalable inference, outperforming Transformers in both performance and efficiency.

### Quantitative analysis of CpG methylation effects on transcription activity

DNA methylation plays a pivotal role in regulating gene expression; however, inferring gene expression from single-cell DNA methylation data remains a major challenge. The following shows scDNAm-GPT’s potential to address this challenge. In the CRC dataset, there are approximately 600 cells with paired single-cell DNA methylation (scDNAm) and single-cell RNA-seq (scRNA-seq) data in the four lesions—primary tumor (PT), lymph node metastasis (LN), liver metastasis (ML), and posttreatment liver metastasis (MP). We split the data into a 7:3 ratio for training and testing. The scDNAm-GPT model was finetuned to predict scRNA-seq expression profiles from single-cell whole-genome methylation information. Further, to test whether the model learns generalizable biological features beyond data memorization, we applied zero-shot transfer learning directly using the expression prediction weights to assess the effect of demethylation at each CpG site on expression levels. This was subsequently validated using other omics data (H3K27ac ChIP-seq and Hi-C/HiChIP) from accessible publications and datasets that the model had never seen before (Fig. 3A).

**Fig. 3.**
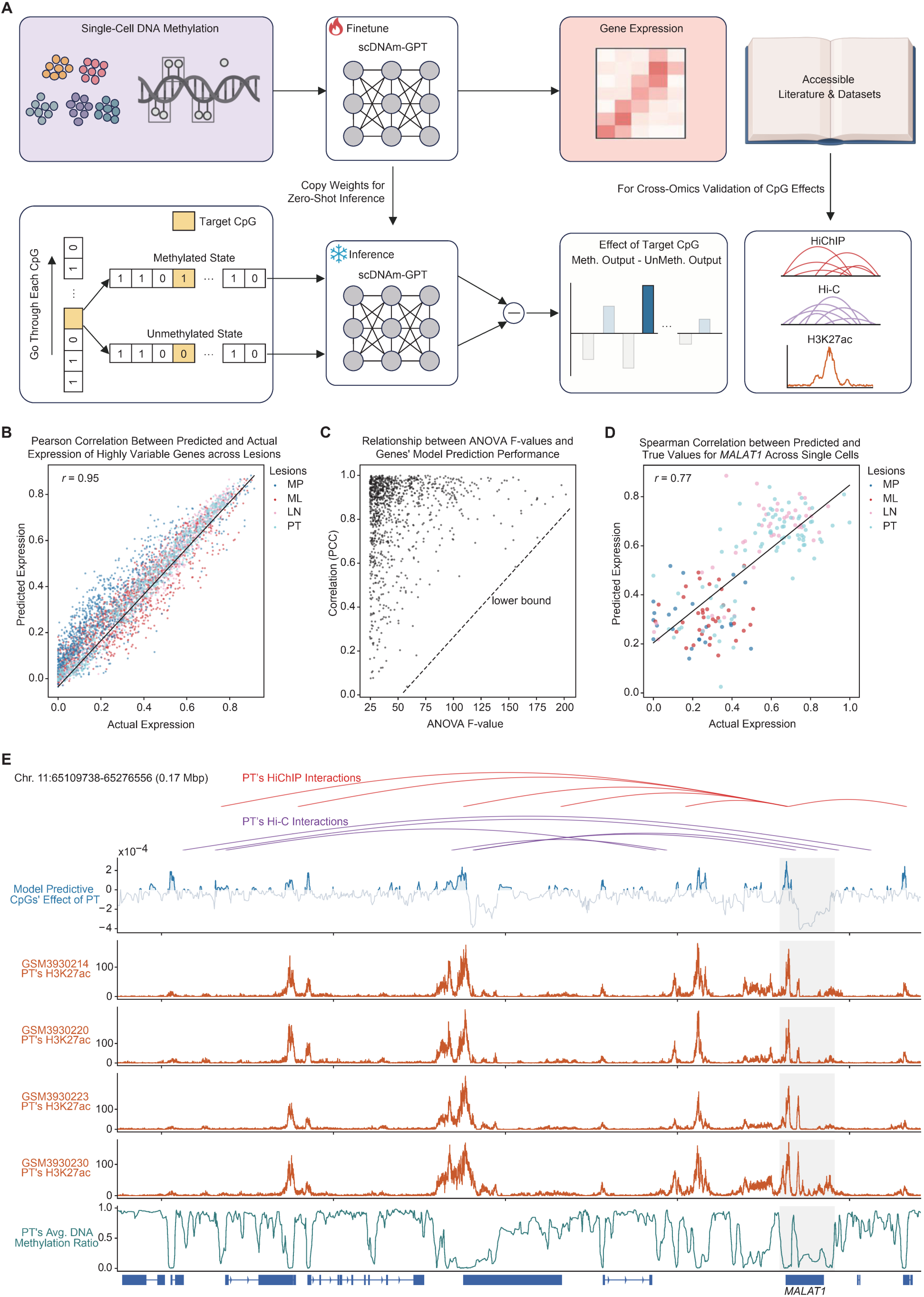
CpG’s effects on gene expression inferred by scDNAm-GPT. (**A**) Schematic of the framework for predicting gene expression from CpG methylation features and quantifying CpG’s effects on gene expression at single-cell resolution. (**B**) Scatterplot illustrating the correlation between actual and scDNAm-GPT-predicted expression levels of highly variable genes among lesions in the test set. (**C**) Relationship between ANOVA F-values and the prediction performance of scDNAm-GPT for highly variable genes among lesions in the test set. (**D**) Scatterplot showing the actual and scDNAm-GPT-predicted expression levels of the *MALAT1* gene across individual single cells in the test set. (**E**) IGV tracks showing scDNAm-GPT-predicted CpG effects, H3K27ac signals, and DNA methylation ratios around the *MALAT1* locus.

The top 1000 highly variable genes across the four lesions (PT, LN, ML, and MP) were selected as the study subjects based on ANOVA F-values, a metric that measures the statistical variance between groups, with higher values indicating greater differences across the groups. The model achieved a Pearson Correlation Coefficient (PCC) of 0.95 between the predicted cancer lesion type expression and the actual expression for the 1000 genes among lesions in the test set (Fig. 3B). Furthermore, we present the correlation between the predicted values and true values for these 1000 genes across different lesions in the test set and analyze the relationship between these genes’ ANOVA F-values and their prediction performance. The results show that genes with low model prediction correlation are enriched at low ANOVA F-values, but these poorly predicted genes progressively disappear as the F-value increases (see the lower bound in Fig. 3C), indicating that the model exhibits greater learning capacity for highly variable genes. Among these 1000 genes, *MALAT1* is a long non-coding RNA that has garnered significant attention in recent cancer research (*19, 20*). Selecting *MALAT1* for further discussion highlights the research value of scDNAm-GPT in unraveling the regulatory mechanisms of cancer-related gene expression. The correlation between the predicted and true values of *MALAT1* across different single cells in the test set reached 0.77 for scDNAm-GPT (Fig. 3D).

To further explore whether the model captures deeper biological mechanisms, the finetuned *MALAT1* expression prediction weights were subjected to zero-shot transfer to assess the contribution of CpG sites around *MALAT1* to its expression levels. In the test set, CpG sites within a 0.17MB region around *MALAT1* in primary tumor cells (PT) were selected for analysis. For each CpG site, both unmethylated and methylated states were input to the model to obtain the predicted *MALAT1* expression levels. The difference between the predicted *MALAT1* expression levels under the unmethylated and methylated states was then calculated and used as the predicted impact of the CpG site on *MALAT1* expression levels (Fig. 3A). The model-predicted effect of each CpG site across all PT cells was averaged and displayed in Fig. 3E as “Model Predicted CpGs’ Effect of PT.”

To assess the interpretability of scDNAm-GPT predictions for CpG effects on gene expression, we included H3K27ac and HiChIP/HiC data derived from CRC. Hi-C reveals 3D chromatin interactions regulating gene expression, HiChIP pinpoints protein-anchored contacts at regulatory loci, and H3K27ac marks active enhancers and promoters, reflecting transcriptional activity. These datasets come from distinct sources (*21*), highlighting the generalizability of the analysis results.

The model-predicted CpGs’ effect on *MALAT1* expression curve closely mirrors the H3K27ac signals, with high-impact sites overlapping chromatin interaction regions identified by HiChIP and Hi-C (Fig. 3E). These results indicate that the model not only fits the data accurately but also captures the underlying regulatory architecture, uncovering the interplay between DNA methylation, transcription, and 3D chromatin organization. It demonstrates the scDNAm-GPT’s ability to understand and model gene regulation at a deeper, mechanistic level.

### Trajectory inference for single-cell DNA methylation data using scDNAm-GPT

The inherently asynchronous states of cells at a given time point enable the application of trajectory inference algorithms to reconstruct dynamic cellular transitions during development and disease progression using single-cell data. To evaluate the capability of scDNAm-GPT in inferring cellular trajectories from DNA methylation profiles, we utilized scWGBS datasets encompassing multiple stages of early embryonic development in both human (*17*) and mouse (*18*). The human dataset includes 280 cells spanning developmental stages from the oocyte to the blastocyst (Fig. 4A). The mouse dataset includes 240 cells spanning developmental stages from the oocyte to the blastocyst. Across both human and mouse datasets, scDNAm-GPT outperforms traditional methods based on DNA methylation ratios, achieving higher concordance with development stages as measured by Kendall’s tau and Spearman’s rho (Fig. 4B). We aggregated the embeddings learned by scDNAm-GPT across all layers and applied diffusion pseudotime analysis to the resulting representations. The model accurately annotated cell stages and successfully reconstructed the developmental trajectory from the zygote to the inner cell mass (ICM) and trophectoderm (TE), in strong concordance with established embryonic progression (Fig. 4C-4E). Similar performance was observed on the mouse dataset (Fig. 4F-4H), further demonstrating the robustness and cross-species generalizability of scDNAm-GPT. Notably, the model was not provided with any temporal annotations; the inferred ordering emerged solely from static DNA methylation profiles following finetuning on cell type supervision. This highlights the model’s ability to capture temporal dynamics from static epigenomic snapshots.

**Fig. 4.**
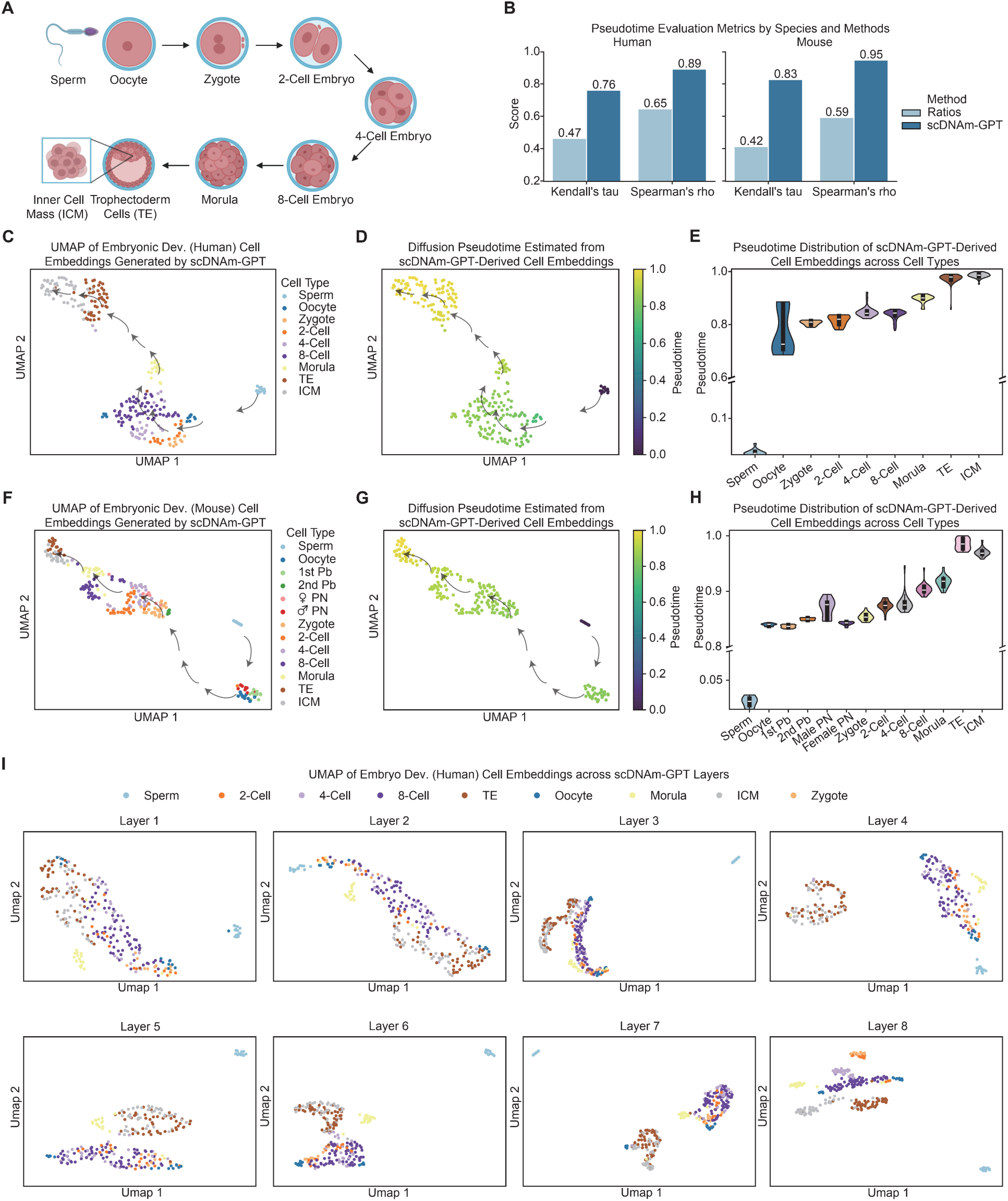
Representation dynamics and pseudotime inference during mammalian early embryo development. (**A**) Schematic of human early embryo development process. (**B**) Kendall’s τ and Spearman’s ρ correlations between predicted pseudotime (inferred from DNA methylation ratios or scDNAm-GPT) and actual pseudotime in human and mouse early embryonic development datasets. (**C-E**) Pseudotime inference for human early embryo development based on aggregated embeddings across all model layers, projected into 2D and colored by inferred pseudotime, revealing a clear developmental trajectory. (**F-H**) Pseudotime inference for mouse early embryo development based on aggregated embeddings across all model layers, projected into 2D and colored by inferred pseudotime, revealing a clear developmental trajectory. (**I**) UMAP visualizations of scDNAm-GPT embeddings from different layers for human early embryo development. Different layers capture distinct types of information, with some layers emphasizing cell-type separation (classification) and others reflecting developmental continuity (pseudotime).

To elucidate how scDNAm-GPT infers dynamic DNA methylation processes, we analyzed the cell embeddings extracted from each layer of the model finetuned on the human dataset. We employed UMAP to visualize the cell representations from all eight layers of scDNAm-GPT, thereby illustrating the progression of feature abstraction and the distinct information captured at each layer (Fig. 4I). In the shallow layers (e.g., layers 1 to 3), only the most pronounced differences were captured—such as the separation of sperm cells from all other cell types—while finer distinctions among more closely related developmental stages were largely omitted, resulting in the mixing of these stages in the embedding space. In contrast, the deeper layers (e.g., layers 4 to 7) captured more nuanced differences, enabling the separation of late-stages (e.g., ICM and TE stages) from early-stages (e.g., Oocyte, Zygote, and Morula). At the eighth layer, each stage forms a compact and well-defined cluster, indicating that the deepest layer encodes highly stage-specific DNA methylation signatures. Similar results were observed on the mouse dataset (Fig. S3). These observations suggest a progressive refinement of developmental trajectories across the model’s hierarchical layers. The results demonstrate that scDNAm-GPT progressively captures both broad and subtle variations in single-cell DNA methylation, enabling a comprehensive representation of epigenetic dynamics with particular sensitivity to temporal regulation.

### The biological interpretability of the context learned by scDNAm-GPT

To investigate how scDNAm-GPT encodes information at the token level, we analyzed the contextual representations learned across its layers. For each layer, we examined the top 1% of genomic regions with the highest attention scores. Analysis of attention distributions revealed that CpG sites with the top 1% highest attention scores for each class exhibited minimal overlap in the early layers (layers 1-4), with Jaccard indices consistently below 0.05. In contrast, the Jaccard index gradually increased from layers 5 to 8, indicating a transition from class-specific to more generalized representations (Fig. 5A). This hierarchical trend suggests that scDNAm-GPT captures progressively complex features of CpG sites, evolving from low-level, localized patterns in shallow layers to high-level, abstract representations in deeper layers. Moreover, the distribution of DNA methylation ratios among the top 1% high-attention regions in layer 8 was notably diverse across classes (Fig. 5B), highlighting the model’s capacity to integrate a broad spectrum of epigenetic information.

**Fig. 5.**
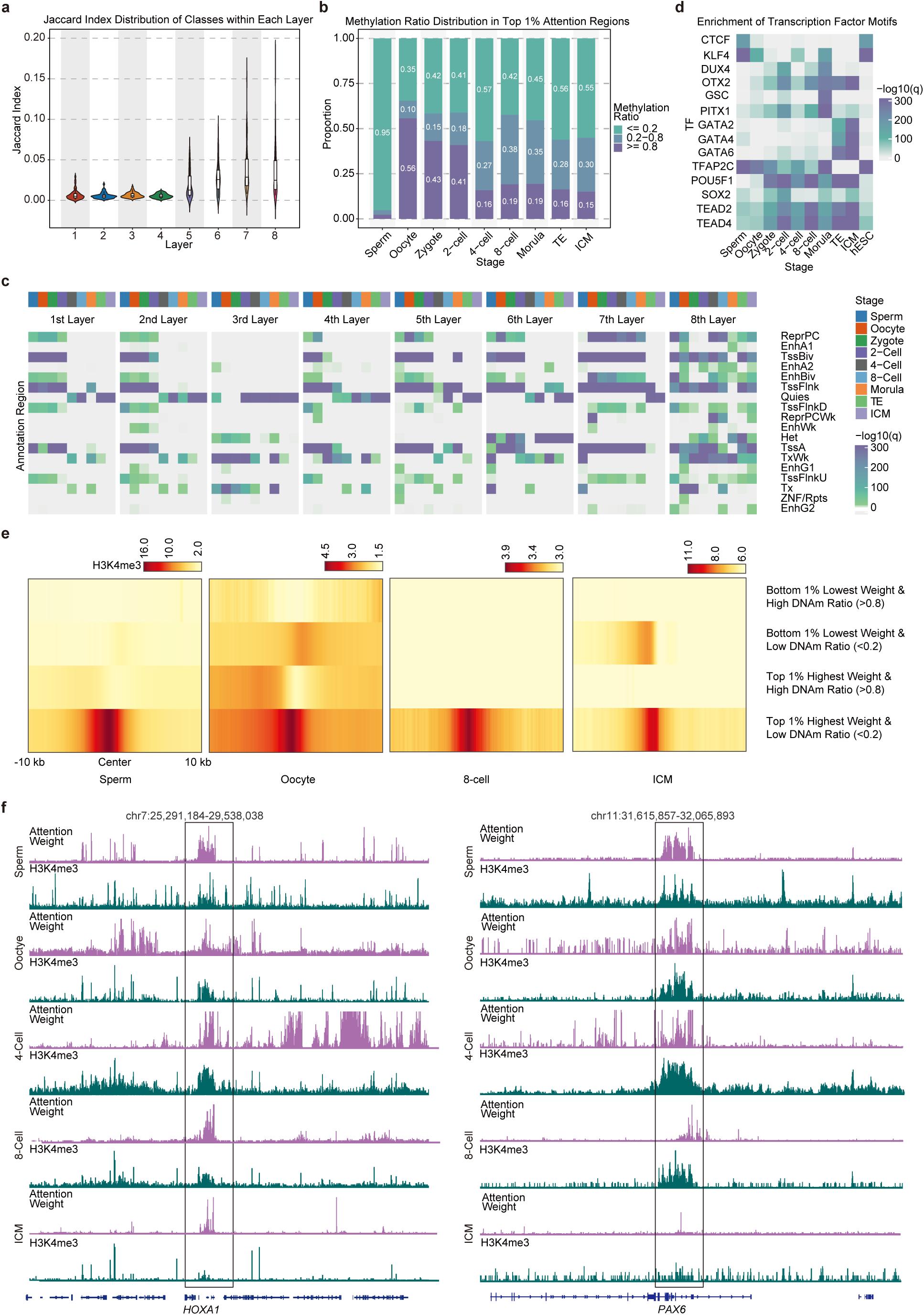
Hierarchical token-level representations and regulatory relevance of scDNAm-GPT attention. (**A**) Jaccard indices of the top 1% high-attention CpG sites across cell classes at each layer. (**B**) Distribution of DNA methylation ratios for the top 1% high-attention CpGs at layer 8, stratified by cell class. (**C**) Chromatin state enrichment analysis of the top 1% high-attention CpGs across layers and development stages. (**D**) Enrichment of transcription factor motifs within the top 1% high-attention regions at layer 8 across development stages. (**E**) Heatmap showing the DNA methylation ratio patterns around the top 1% high-attention and bottom 1% low-attention CpG sites, stratified by DNA methylation ratio at layer 8. (**F**) IGV browser snapshots showing attention scores and H3K4me3 ChIP-seq signals at the *HOXA1* and *PAX6* locus.

To further characterize the contextual learning of the model, we performed chromatin state enrichment analysis on the top 1% attention-scoring regions. Distinct enrichment patterns were observed across layers. Layers 1 to 4 showed significant enrichment in transcription start site (TSS) regions while being depleted for enhancer and repeat-associated regions. In contrast, layers 5 to 8 displayed broader enrichment profiles, including TSSs, enhancers, and repetitive elements. Notably, layer 8 exhibited comprehensive coverage of diverse regulatory elements across different cell types (Fig. 5C). This progressive expansion in chromatin context suggests that scDNAm-GPT learns regulatory features in a hierarchical manner, moving from narrow, localized signals to a diverse and integrated view of the epigenomic landscape.

We further conducted transcription factor (TF) binding motif enrichment analysis using the JASPAR database. This analysis revealed dynamic and stage-specific motif enrichment within the top 1% high-attention regions identified by layer 8 (Fig. 5D). For example, motifs corresponding to inner cell mass (ICM)-specific TFs such as GATA2, GATA4, and GATA6 were significantly enriched in ICM-associated regions. In embryonic stem cell (hESC) stages, motifs for CTCF and TFAP2C were prominently enriched. At the 8-cell stage, motifs corresponding to OTX2 and PITX1 were significantly overrepresented. These findings are consistent with previously reported stage-specific TF activity profiles (*22*), supporting the biological relevance of the model’s attention-based prioritization.

To further explore the epigenomic context of the model’s high-attention regions, we integrated histone modification data. We observed that regions with top 1% attention scores and low DNA methylation levels (<0.2) were significantly enriched for the active histone mark H3K4me3. In contrast, high-attention regions with high DNA methylation ratios (>0.8) showed marked depletion of H3K4me3 signals. Notably, regions within the lowest 1% attention scores—regardless of DNA methylation status—did not exhibit any significant H3K4me3 enrichment (Fig. 5E). These results suggest that scDNAm-GPT preferentially attends to regulatory regions marked by active chromatin features, particularly within hypomethylated contexts. For example, at the *HOXA1* locus, regions with high attention scores aligned with areas of enriched H3K4me3 signal across various early embryonic stages (Fig. 5F), in agreement with previous findings (*23*). A similar pattern was also observed in the *PAX6* locus, which is also consistent with previous findings (*23*).

Collectively, these analyses demonstrate that scDNAm-GPT not only learns meaningful sequence-level and methylation-level representations but also captures biologically interpretable and hierarchically structured features of the epigenomic landscape.

### Reference-Free deconvolution of heterogeneous bulk DNA methylation profiles by scDNAm-GPT

Accurate inference of the cellular composition of bulk DNA methylation samples—such as those derived from cell-free DNA—is a critical task known as cell type deconvolution (Fig. 6A-B). Traditional deconvolution methods typically rely on predefined reference profiles for specific cell types, which are often limited by the difficulty of obtaining comprehensive and representative profiles, the lack of generalizability across tissues and conditions, and their inability to capture the extensive heterogeneity within individual cell types. To address these limitations, we propose a reference-free, end-to-end deconvolution strategy leveraging the pretrained scDNAm-GPT model.

**Fig. 6.**
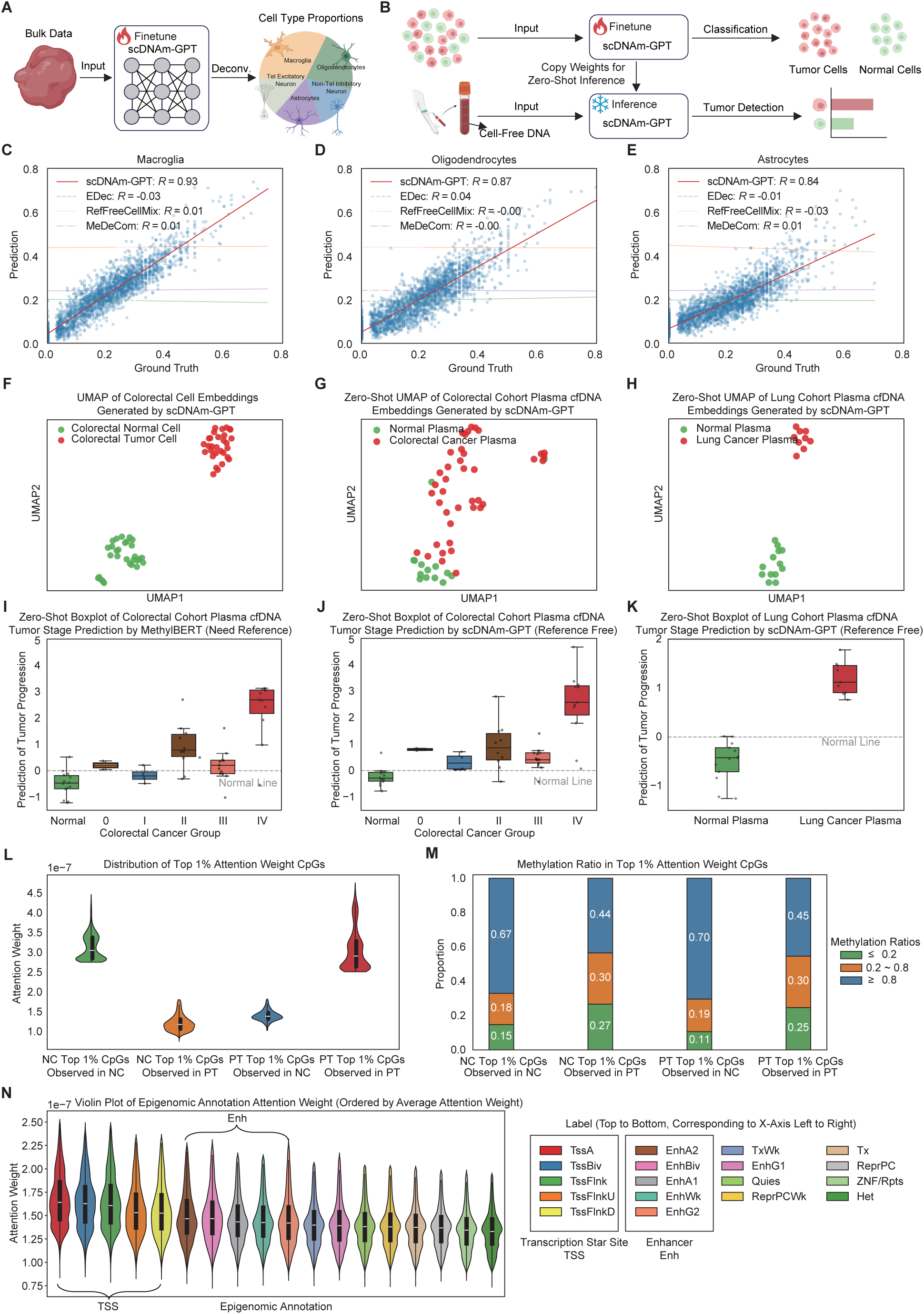
Reference-free deconvolution and cfDNA-based cancer detection using scDNAm-GPT. (**A**) Schematic of reference-free deconvolution strategy. (**B**) Schematic illustration of cfDNA-based cancer detection using single-cell DNA methylome data as input. (**C-E**) Predicted vs. true average cell-type (macroglia, Oligodendrocytes, and Astrocytes) proportions in simulated bulk methylation samples across different neuronal types (R = 0.84 ∼ 0.93). (**F**) UMAP of cell embeddings of colorectal cells generated by scDNAm-GPT. (**G**) Zero-shot UMAP visualization of colorectal cancer patients based on plasma cfDNA methylation profiles inferred by scDNAm-GPT. (**H**) Zero-shot UMAP visualization of lung cancer patients based on plasma cfDNA methylation profiles inferred by scDNAm-GPT. (**I, J**) Zero-shot boxplot showing colorectal tumor cfDNA fraction from plasma inferred by MethylBERT (need reference) and scDNAm-GPT (reference free). (**K**) Zero-shot boxplot showing lung tumor cfDNA fraction from plasma inferred by scDNAm-GPT. (**L**) Violin plot showing the distribution of the top 1% high-attention CpG sites’ attention weights in NC and PT. (**M**) Histogram showing DNA methylation ratio composition on the top 1% high-attention CpG sites in NC and PT. (**N**) The distribution of attention weights on regulatory elements.

In silico mixtures with known cell type proportions are essential for evaluating deconvolution performance. Previously published methods generated DNA methylation signals for each cell type by sampling from a single randomly selected methylome profile within that cell type (*24*). However, this approach overlooks the inherent variability among samples of the same cell type. We simulated bulk DNA methylation data using single-cell DNA methylation data from five different neuronal cell types (*11*). For each simulation, we randomly sampled between zero and ten cells from each cell type to generate mixtures that preserved both inter-cell-type and intra-cell-type heterogeneity. To better mimic real bulk sequencing conditions, we maintained variations in sequencing depth and coverage across the mixed single-cell samples. These variations reflect technical noise arising from differences in capture efficiency among cells and uneven sequencing depth across genomic loci.

The cell type proportions predicted by scDNAm-GPT showed high concordance with the true proportions across five distinct neuronal subtypes, achieving an average Pearson correlation coefficient of 0.88 (Fig. 6C-6E). In contrast, other reference-free deconvolution methods—RefFreeCellMix (*25*), EDec (*26*), and MeDeCom (*27*)—were unable to accurately infer cell type compositions (Fig. S4). This discrepancy may be attributed to the subtle DNA methylation differences among the five neuronal subtypes, which are not readily captured by previous approaches primarily designed to distinguish cell types across divergent lineages. These findings highlight the superior capability of scDNAm-GPT in resolving fine-grained cell type compositions from DNA methylation data, particularly within closely related cell subtypes, such as neuronal subtypes.

To further assess the model’s ability to detect disease-associated DNA methylation patterns, we finetuned scDNAm-GPT on a binary classification task using single-cell whole-genome bisulfite sequencing (scWGBS) data from healthy colorectal cells and primary colorectal tumor cells. The finetuned embeddings exhibited clearly separated clusters corresponding to normal and tumor cells, demonstrating that scDNAm-GPT effectively captures tumor-specific DNA methylation signatures (Fig. 6F). We then conducted a zero-shot evaluation by directly applying the tissue-finetuned model to circulating cell-free DNA (cfDNA) from peripheral blood, colon cancer (*28*), and lung cancer (*29*) patients, without any additional training on cfDNA data. Remarkably, the model robustly distinguished cfDNA from healthy individuals and colorectal cancer patients across distinct clinical stages, as well as from lung cancer patients (Fig. 6G, 6H). We next benchmarked scDNAm-GPT against MethylBERT (*3*), a state-of-the-art reference-based cfDNA deconvolution method. While scDNAm-GPT reliably separated cfDNA from healthy individuals and cancer patients, MethylBERT failed to clearly distinguish several stages I and III cases (Fig. 6I-6K). Notably, we introduce a framework that finetunes scDNAm-GPT using a limited number of scWGBS or bulk DNA methylation samples, eliminates manual feature selection and reference construction, and achieves improved deconvolution performance with enhanced clinical relevance. These results highlight strong generalization capability of scDNAm-GPT and its superior ability to robustly capture tumor-specific DNA methylation features even under the high background noise characteristic of cfDNA.

Importantly, we found that the top 1% of high attention weights differed significantly between healthy and primary tumor cells (Fig. 6L). Moreover, the DNA methylation ratio distributions within these high-attention regions also showed clear differences, suggesting that scDNAm-GPT captures DNA methylation features in a context-specific manner (Fig. 6M). We observed that the attention weights assigned by scDNAm-GPT exhibit distinct genomic region preferences. Specifically, attention is highest around transcription start sites (TSS), suggesting the model prioritizes regulatory signals closely linked to gene activation. Enhancer regions also show elevated attention, reflecting their role in gene regulation through distal interactions. In contrast, repressive regions and heterochromatin display relatively lower attention weights, consistent with their association with transcriptional silencing and reduced regulatory activity (Fig. 6N). These patterns indicate that scDNAm-GPT effectively captures biologically relevant features of the DNA methylation landscape and assigns importance in a manner aligned with known epigenomic functional elements.

## Discussion

We introduce scDNAm-GPT, the first universal large language model designed for single-cell DNA methylation analysis. Our model can process up to 20 million CpG sites simultaneously and integrate over 80 million nucleotides of surrounding sequence context, opening the door to a new era of genome-wide single-cell omics modeling. It is built upon five key innovations that distinguish it from previous approaches: i) CpG-centered 6-mer tokenization, which captures both the methylation state and the surrounding sequence context at each CpG site; ii) Mamba’s Selective State Space Models (SSMs) (*8*), enabling efficient processing of high-dimensional, sparse data; iii) the use of cross-attention heads, which integrate and align local CpG features with broader sequence context to capture long-range dependencies and subtle epigenetic signals; iv) an innovative fuzzy token input design, specifically tailored for DNA methylation, that allows seamless compatibility with both single-cell and bulk DNA methylation data; and v) a novel next fuzzy token prediction pretraining strategy, which enhances the model’s ability to learn probabilistic methylation patterns and subtle epigenetic variations. By leveraging these innovations and pretraining on over one million single-cell DNA methylation profiles, scDNAm-GPT effectively captures complex interactions among CpG sites, providing a robust tool for downstream analyses and generating biologically meaningful embeddings of epigenetic states.

Due to the sparsity and high dimensionality of scWGBS data, clustering and cell type prediction remain significant challenges. Traditional methods mitigate sparsity by averaging DNA methylation ratios over 100,000 bp bins; however, this approach also obscures informative DNA methylation signals and leads to inconsistent clustering results. In contrast, scDNAm-GPT achieves a clustering accuracy of 96% and effectively generates meaningful clustering results that align with cell types, even for unseen data. Notably, regions with higher attention weights correspond to low DNA methylation ratios and are marked by active histone modifications, underscoring the interpretability of our model. The sizes of regions with higher attention scores range from hundreds to tens of thousands of base pairs, demonstrating the model’s superior resolution in identifying biologically meaningful DNA methylation regions.

Beyond clustering and cell type prediction, scDNAm-GPT demonstrates promising capabilities in reconstructing cell differentiation trajectories. This feature provides novel insights into epigenetic dynamics during development and lineage commitment, positioning our model as a powerful tool for investigating cell fate decisions and the underlying mechanisms of epigenetic regulation. Furthermore, preliminary results suggest that scDNAm-GPT can competitively deconvolve cell type compositions from cell-free DNA methylation data in a reference-free manner, addressing a challenge that remains difficult for traditional reference-based approaches.

scDNAm-GPT has certain limitations. First, although the pretraining dataset encompassed diverse tissues and cell types, it was heavily weighted toward brain data, potentially limiting the model’s ability to fully and equally capture the epigenetic features of other organs and disease contexts. Expanding the training dataset to incorporate scWGBS profiles from a broader range of organs, diseases, and cell types will be critical to enhancing the model’s generalizability and predictive power in future applications. Additionally, scDNAm-GPT currently does not integrate other omics data, such as scRNA-seq, scHi-C, or scCUT&Tag, which could enhance its applicability across various tasks. Furthermore, incorporating metadata such as individual patient information, clinical imaging, and physiological parameters (e.g., blood pressure) could significantly enhance scDNAm-GPT’s ability to predict clinical status at an individual level.

Time-series and perturbation data are needed to enable causal relationships to be learned and cell behavior to be inferred in response to DNA methylation alterations. As a foundation model, scDNAm-GPT is positioned to accelerate the discovery of single-cell epigenetic dynamics, advancing biomedical research.

## Methods

### 1. Model

#### 1.1. scWGBS Tokenizer

The scWGBS tokenizer is a critical component of scDNAm-GPT, as it transforms raw genomic data into a format suitable for processing by the model. Given that scWGBS data consists of methylation levels at each CpG site across large genomic regions, the tokenizer’s primary role is to encode these methylation values into a tokenized sequence that preserves both the spatial and biological characteristics of the input data.

In contrast to traditional NLP tokenization, where words are split into subwords or characters, the scWGBS tokenizer is designed to handle the unique challenges posed by biological data, particularly the representation of DNA sequences and methylation patterns. Each CpG site is treated as a “token,” and its methylation state is encoded in a way that retains the crucial epigenetic information.

The core idea behind the scWGBS tokenizer is to treat DNA sequences as strings of “CpG words,” where each CpG site is a token. This tokenizer design incorporates both methylated and unmethylated states as part of the token representation, ensuring that methylation patterns are preserved for downstream processing.

To achieve this, each CpG site is encoded as a binary token that represents its methylation state: - A methylated CpG is assigned a token value of 1, - An unmethylated CpG is assigned a token value of 0.

In addition to binary methylation states, we incorporate flanking nucleotide context to account for the surrounding DNA sequence, which can have biological significance in regulating methylation patterns. This allows the tokenizer to generate more informative representations for each CpG site.

Thus, each token is a combination of the methylation state and its local DNA context, forming a sequence of tokens that represent the entire genomic region.

The tokenization process can be broken down into the following steps:

1. DNA Sequence Splitting: The genomic data is split into windows of CpG sites, where each window corresponds to a continuous region of the genome (e.g., a 10kb region). Each window contains a sequence of CpG sites, and the surrounding nucleotides are encoded as part of the context.
2. Encoding Methylation States: For each CpG site in the window, its methylation state is encoded as a binary value (0 or 1). Additionally, the surrounding nucleotides (e.g., ± 3 bases around each CpG site) are incorporated to form a context-aware token. This results in a k-mer token representation, where each CpG is accompanied by its local sequence context.
3. Sequence Assembly: The tokenized sequences are then assembled into a sequence of tokens, where each token represents a CpG site along with its methylation state and surrounding context. The final sequence represents the entire window of genomic data.

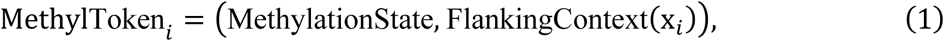

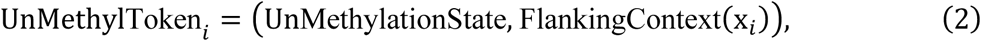 Where *x_i_* is the *i*-th CpG site in the genomic window, and the MethylationState and the UnMethylationState refer to MG and CG, respectively, while the FlankingContext represents the surrounding nucleotides.
4. *[BOS]* and *[SEP]* Token Insertion: A special *[BOS]* token is inserted at the beginning of each window to signal the start of a new sequence, as well as to serve as the query in the cross-attention mechanism, which aggregates global context from the entire sequence.

Once the tokenization process is complete, the resulting sequence of tokens can be fed into the Mamba Backbone for further processing. The *[SEP]* token plays a crucial role in generating the query vector for the cross-attention mechanism, as described in Section A.1.3. This integration of tokenized genomic data with the cross-attention mechanism allows scDNAm-GPT to capture both the methylation state of individual CpG sites and their relationships with distant sites across the genome.

For example, consider a window of CpG sites:

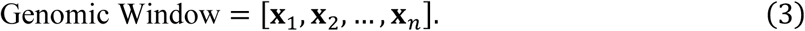

The tokenizer processes this window by encoding each CpG site **x*_i_*** as a token that includes its methylation state and flanking nucleotide context. The resulting sequence is then represented as:

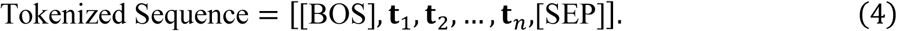

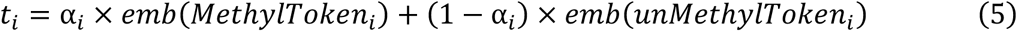

This tokenized sequence is then passed through the Mamba Backbone, where the *[SEP]* token generates the query vector for the cross-attention mechanism, which attends to each CpG site (represented by the tokens **t*_i_*** based on their methylation states and contextual relationships.

The scWGBS tokenizer offers several advantages for modeling DNA methylation data:

1. Preserving Methylation Information: By encoding both methylation states and surrounding nucleotide contexts, the tokenizer ensures that key epigenetic information is preserved for downstream processing.
2. Contextualized Tokenization: The inclusion of flanking nucleotides in the tokenization process allows the model to learn not only the methylation state of each CpG site but also the surrounding sequence context that may influence methylation patterns.
3. Scalability: The tokenizer’s ability to efficiently process large genomic windows and generate contextually rich tokens enables scDNAm-GPT to scale to millions of CpG sites, making it suitable for analyzing large-scale genomic data.

The scWGBS tokenizer is specifically designed to capture biologically relevant features of the genome. The methylation status of each CpG site is crucial for understanding gene regulation, and the flanking nucleotide context helps reveal potential regulatory motifs that influence methylation patterns. By integrating these features into the tokenization process, we ensure that the model can learn meaningful biological patterns from the data.

#### 1.2. Mamba Backbone

Mamba is a state-of-the-art sequence modeling framework designed for efficient processing of ultra-long sequences. Unlike traditional transformer-based architectures, which suffer from quadratic complexity in self-attention, Mamba leverages Selective State Space Models (SSMs) (*7*) to achieve linear time complexity while maintaining strong representation capabilities.

The motivation for using Mamba in scDNAm-GPT stems from the inherent challenges of single-cell whole-genome bisulfite sequencing (scWGBS) data. Each cell’s methylation profile consists of millions of CpG sites, forming an ultra-long sequence that requires efficient modeling of both local CpG interactions and long-range dependencies. Mamba’s ability to capture global dependencies without the prohibitive computational costs of transformers makes it a very natural fit for this problem.

Long genomic sequences present unique challenges in computational biology (*30, 31*) Methylation patterns at distant CpG sites often exhibit correlations due to chromatin organization, regulatory regions, and epigenetic inheritance. Traditional deep learning models, including recurrent neural networks (RNNs) and CNNs, struggle with these dependencies due to limited receptive fields or memory constraints.

Mamba addresses these limitations through a continuous-time selective state space model (SSM), which allows for efficient and memory-friendly long-sequence modeling. The core state update equation is:

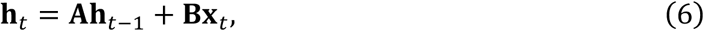

where: **h*_t_*** is the hidden state at time step *t*, **A** is a learned transition matrix controlling sequence memory, **B** is an input transformation matrix, **x*_t_*** is the input at time step *t*

Unlike transformers, which require explicit attention over all previous positions, Mamba’s state-space model implicitly propagates long-range dependencies without explicit considering the token-by-token interactions, significantly reducing computational overhead.

The Mamba Backbone in scDNAm-GPT consists of a deep stack of **Mamba Blocks**, each designed to refine feature representations at increasing levels of abstraction. Each block operates as follows:

1. Convolutional Input Projection: The input sequence X is first processed by a depth-wise convolutional layer to extract local CpG patterns:

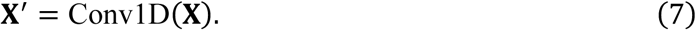
2. State-Space Model (SSM) Evolution: Each token is passed through a continuous-time state space layer, updating the internal sequence representation:

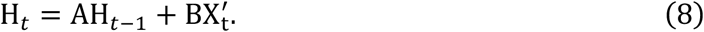
3. Selective Gating Mechanism: A sigmoid gating function dynamically modulates the influence of new information:

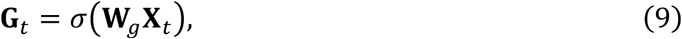

where *σ* is the sigmoid activation and **W*_g_*** is a learned weight matrix.
4. Linear Projection and Normalization: The output is projected to a new feature space and normalized:

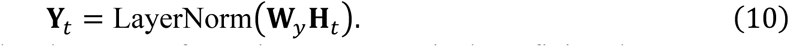

Each Mamba Block stacks these transformations, progressively refining long-sequence representations.

Network Configuration and Parameters

The Mamba Backbone in scDNAm-GPT consists of 8 stacked Mamba Blocks, leading to a total parameter count of approximately 1M parameters. This configuration balances computational efficiency with modeling capacity.

The architecture can be expressed as a sequential composition:

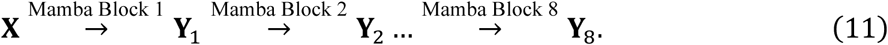

This deep stacking mechanism enables the model to capture both short-range CpG patterns and long-range regulatory dependencies in scWGBS data. The Mamba Block computations are optimized using CUDA kernels for efficient parallelization, allowing scDNAm-GPT to process ultra-long sequences efficiently.

#### 1.3. Cross Attention Head

Cross-attention is a powerful mechanism that allows a model to focus on relevant parts of the input sequence while processing a different sequence. In traditional attention mechanisms, such as in the Transformer model, a query vector interacts with all keys and values within the same sequence. However, in cross-attention, the query vector is derived from one sequence, and it attends to another sequence’s keys and values. This approach is especially useful when the relationships between two different data representations are essential, such as in multi-modal tasks or when integrating sparse information like DNA methylation patterns.

In our scDNAm-GPT framework, we leverage cross-attention to capture long-range dependencies and contextualize CpG site interactions across the entire genome. Specifically, we use a *[SEP]* token to generate query, key, and value representations (denoted as **Q, K, V**) for the cross-attention mechanism. The *[SEP]* token’s final hidden state serves as the query, while each CpG site’s hidden state is used to compute the keys and values.

The mechanism of the employed cross-attention is clarified as follows:

Given an input sequence **X = [x_1_, x_2_, …, X*_n_*]**, where **x*_i_*** represents the hidden state corresponding to the *i*-th token in the sequence, cross-attention computes the interactions between the query **Q** and the keys **K**, along with their corresponding values **V**.

The cross-attention operation can be described as:

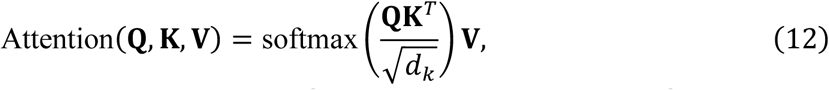

where: 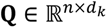 is the query matrix, 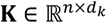 is the key matrix, 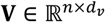 is the value matrix, *d_k_* is the dimension of the key vectors.

The softmax function ensures that the attention weights sum to 1, providing a probabilistic distribution over the values **V** based on the relevance of the queries to the keys.

In scDNAm-GPT, we utilize cross-attention to capture the intricate dependencies between CpG sites across the entire genome, focusing on biologically meaningful interactions. To achieve this, we employ the *[SEP]* token in the following way:

1. The *[SEP]* token’s final hidden state is used as the query **Q** for cross-attention. This token is specially designed to aggregate global contextual information from the entire sequence.
2. Each CpG site’s hidden state, generated from the Mamba Backbone, serves as both the key **K** and the value **V**.

Thus, for each position in the sequence (representing a CpG site), the attention weights are computed between the *[SEP]* token’s query and the CpG site’s key. This allows the *[SEP]* token to attend to the relevant CpG sites and capture their relationships in the final representation.

The cross-attention operation for a single CpG site can be described as:

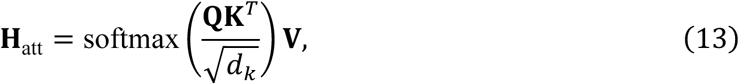

where: **H**_att_ represents the attention-weighted hidden state for each CpG site after the cross-attention computation, **Q** is the hidden state of the *[SEP]* token, **K**, and **V** are the hidden states of each CpG site.

This attention-weighted hidden state **H**_att_ is then used for further processing in downstream layers, where it contributes to the final genomic sequence representation.

The use of cross-attention in scDNAm-GPT offers several benefits for modeling genomic data:

1. Capturing Long-Range Dependencies: Cross-attention allows the *[SEP]* token to attend to distant CpG sites, which is essential for modeling the long-range interactions that occur in DNA methylation patterns. These long-range dependencies often play crucial roles in gene regulation and epigenetic modifications.
2. Contextualizing CpG Interactions: By attending to the entire genomic sequence, the cross-attention mechanism helps the model contextualize the interaction between CpG sites, ensuring that biologically relevant correlations are learned.
3. Efficient Representation Learning: The ability of the *[SEP]* token to serve as a global query ensures that even sparse methylation data can be processed effectively, enabling the model to capture meaningful patterns from incomplete or sparse data.

The cross-attention mechanism is particularly effective in processing large-scale genomic datasets, such as those in scDNAm-GPT, where each input sequence can span millions of CpG sites. By using the *[SEP]* token to generate the query vector, we avoid the need for expensive pairwise attention calculations across all tokens, enabling the model to efficiently capture global dependencies while maintaining computational feasibility.

In scDNAm-GPT, the cross-attention mechanism is integrated within the architecture to handle ultra-long sequences efficiently. By applying attention selectively through the *[SEP]* token query, the model can scale to handle large genomic sequences with millions of CpG sites while still extracting biologically meaningful relationships between distant CpG sites.

### 2. Pretraining Loss Function

Given:

- *X*: the sequence of input tokens (CpG sites and context).
- *y_unmethy_*: labels for unmethylated CpG sites.
- *y_methy_*: labels for methylated CpG sites.
- *α_methy_*: methylation ratio (replaced *r* with *α*).
- *ŷ*: predicted logits (output of the model).

1. Shifted logits and labels for causal language modeling

- ŷ_shifted_ = ŷ[1:]
- y_unmethy,shifted_ = y_unmethy_[1:]
- y_methy,shifted_ = y_methy_[1:]
- α_methy,shifted_ = α_methy_[1:]
2. CrossEntropy loss:The model computes the cross-entropy loss for both methylated and unmethylated CpG sites, weighted by their respective methylation ratios (*α)*:

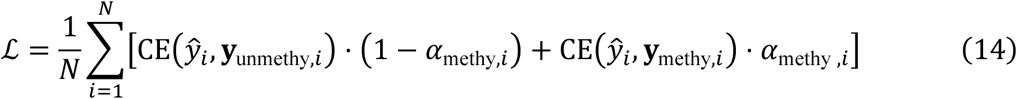 Where *CE* is the cross-entropy function and *N* is the number of CpG sites.
3. Pretraining loss calculation: The pretraining loss is computed by applying a mask to exclude padding tokens and averaging the remaining loss values:

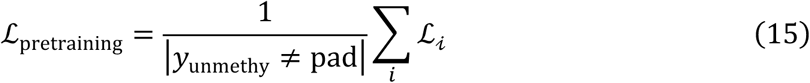
4. Core concept: fuzzy next token prediction:The core idea is to predict the next token (CpG site) given the current state, while considering both the methylation and unmethylation states of CpG sites. The loss is fuzzy in that it accounts for the methylation ratio, α, effectively weighting the prediction for methylated and unmethylated CpGs differently, based on their respective methylation status.

### 3. Hyperparameters

The parameters commonly used include: max_steps (the maximum number of training steps), set to 30,000 steps; eval_strategy and eval_steps, which define the evaluation strategy and frequency, with evaluation occurring every 100 steps; save_steps and logging_steps, indicating that the model is saved and logs are recorded every 100 steps; fp16/8, enabling 16/8-bit floating-point computation to enhance training speed and reduce memory usage; learning_rate set to 3e-05, controlling the step size for model weight updates; lr_scheduler_type, using a linear scheduler to decrease the learning rate gradually; warmup_steps, set to 100 steps, allowing for a gradual increase in learning rate at the start of training; and max_grad_norm, which limits the maximum gradient norm to 10.0 to prevent gradient explosion. Additionally, the batch size (batch_size) is set to 16, meaning 16 samples are used for each training step. These parameter settings collectively ensure efficient and stable training. Additionally, the training was performed on H200 system. With fp16 precision, the maximum sequence length (max_length) that can be processed is 21 million tokens, while with fp8 precision, the system can handle up to 35 million tokens. These configurations ensure efficient, stable, and scalable tuning scDNAm-GPT on large datasets.

### 4. Evaluation Metrics

This section provides detailed definitions and mathematical formulations for the evaluation metrics used throughout this study to assess model performance on various tasks, including classification, language modeling, correlation, and rank-order comparison.

Accuracy (ACC): Accuracy measures the proportion of correctly classified instances over the total number of instances. It is a fundamental metric for classification tasks, such as cell type annotation.

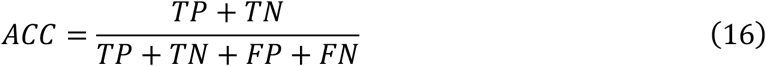

where TP, TN, FP, and FN represent the number of True Positives, True Negatives, False Positives, and False Negatives, respectively.

F1-score: The F1-score is the harmonic mean of Precision and Recall, providing a single metric that balances both concerns. It is particularly useful for evaluating classification performance on datasets with imbalanced class distributions.

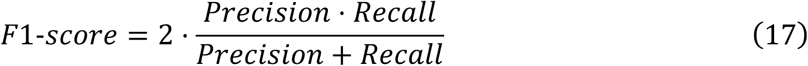

where Precision is the proportion of true positive instances among all instances classified as positive, calculated as:

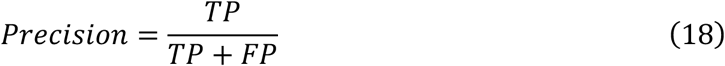

Recall (Sensitivity or True Positive Rate): Recall measures the model’s ability to identify all relevant instances within a dataset. It is the proportion of true positive instances that were correctly classified, and it is particularly important in scenarios where failing to detect a positive case (e.g., a specific cell type) is critical.

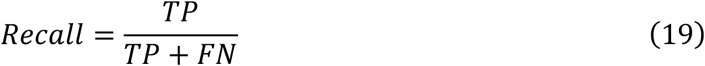

Perplexity (PPL): Perplexity is a standard metric for evaluating the performance of language models. It measures how well a probability model predicts a sample, with a lower value indicating better performance. It is defined as the exponentiated average negative log-likelihood of a sequence.

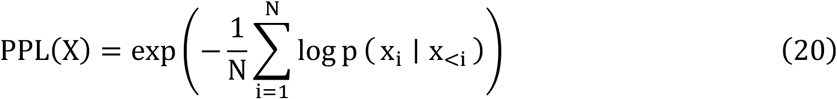

where *X* = (*x*_1_, *x*_2_, … *x_N_*) is a sequence of *N* tokens, and *p*(*x_i_* | *x*_<1_) is the conditional probability of the *i*-th token given the preceding tokens, as assigned by the model.

Kendall’s rank correlation coefficient (*τ*): Kendall’s tau is another non-parametric rank correlation coefficient that measures the ordinal association between two quantities. It assesses the similarity of the ordering of the data when ranked by each of the quantities. This metric is particularly well-suited for comparing predicted cellular pseudotime with actual developmental stages.

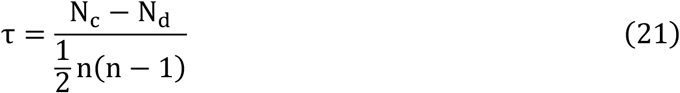

where *n* is the number of items, *N_c_* is the number of concordant pairs (pairs of items ordered in the same way by both rankings), and *N_d_* is the number of discordant pairs (pairs of items ordered differently by the two rankings).

Spearman’s rank correlation coefficient (*ρ*): Spearman’s rho is a non-parametric measure of the monotonic relationship between two variables. It assesses how well the relationship can be described using a monotonic function by calculating the Pearson correlation coefficient on the rank-transformed variables. It is robust to outliers and does not assume a linear relationship.

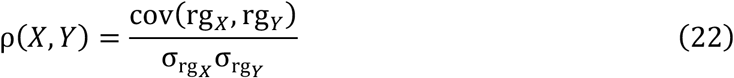

where rg*_X_* and rg*_Y_* are the rank variables corresponding to the original variables *X* and *Y*.

Pearson Correlation Coefficient (PCC): The Pearson Correlation Coefficient measures the linear relationship between two continuous variables. It is used here to assess the concordance between predicted and actual values, such as gene expression levels. Its value ranges from +1 (total positive linear correlation) to −1 (total negative linear correlation), with 0 indicating no linear correlation.

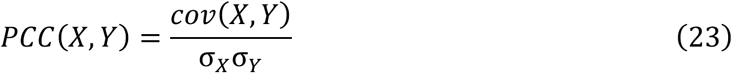

where *X* and *Y* are the two variables, *cov*(*X*, *Y*) is their covariance, and *σ_X_* and *σ_Y_* are their respective standard deviations.

## Code availability

Code and model weights necessary for reproducing the results is available at GitHub (https://github.com/ChaoqiLiang/scDNAm-GPT). The code is released under the MIT License. There are no restrictions on the availability of the data or materials, except where required by materials transfer agreements (MTAs). All data are available in the main text or the supplementary materials.

## Acknowledgments

We thank the technical support from the Data Science Platform of Guangzhou National Laboratory and the Bio-medical Big Data Operating System (Bio-OS). We thank the technical support from the JC STEM Lab of AI for Science and Engineering.

## Funding

National Natural Science Foundation of China (82370148) (JL)

R&D Program of Guangzhou Laboratory (GZNL2023A02003) (JL)

R&D Program of Guangzhou Laboratory (GZNL2023A02011) (JL)

Guangzhou Municipal Science and Technology Bureau, Guangzhou Key

Research and Development Program (2024B03J0046) (JL)

Supported by the grant of State Key Laboratory of Respiratory Disease

(SKLRD-Z-202307) (JL)

The Hong Kong Jockey Club Charities Trust, the Research Grants Council of Hong Kong (Project No. CUHK14213224). (WLO)

## Author contributions

Conceptualization: JL, CQL, PY, HLY

Methodology: CQL, PY, HLY, PZ, WMZ, LB, WLO

Investigation: JL, CQL, PY, HLY, JLS, YNW, YL, YCR, YPJ, RW, JJX, SZZ, LLJ, WQB, XZM, TC

Visualization: PZ, CQL, YL, YNW

Funding acquisition: JL, WMZ, LB, WLO

Project administration: PY

Software: CQL, PZ

Supervision: JL, WMZ, LB, WLO

Writing – original draft: CQL, JL, PY, HLY, PZ

Writing – review & editing: CQL, JL, PY, HLY, PZ

## Supplement

**Figure S1.**
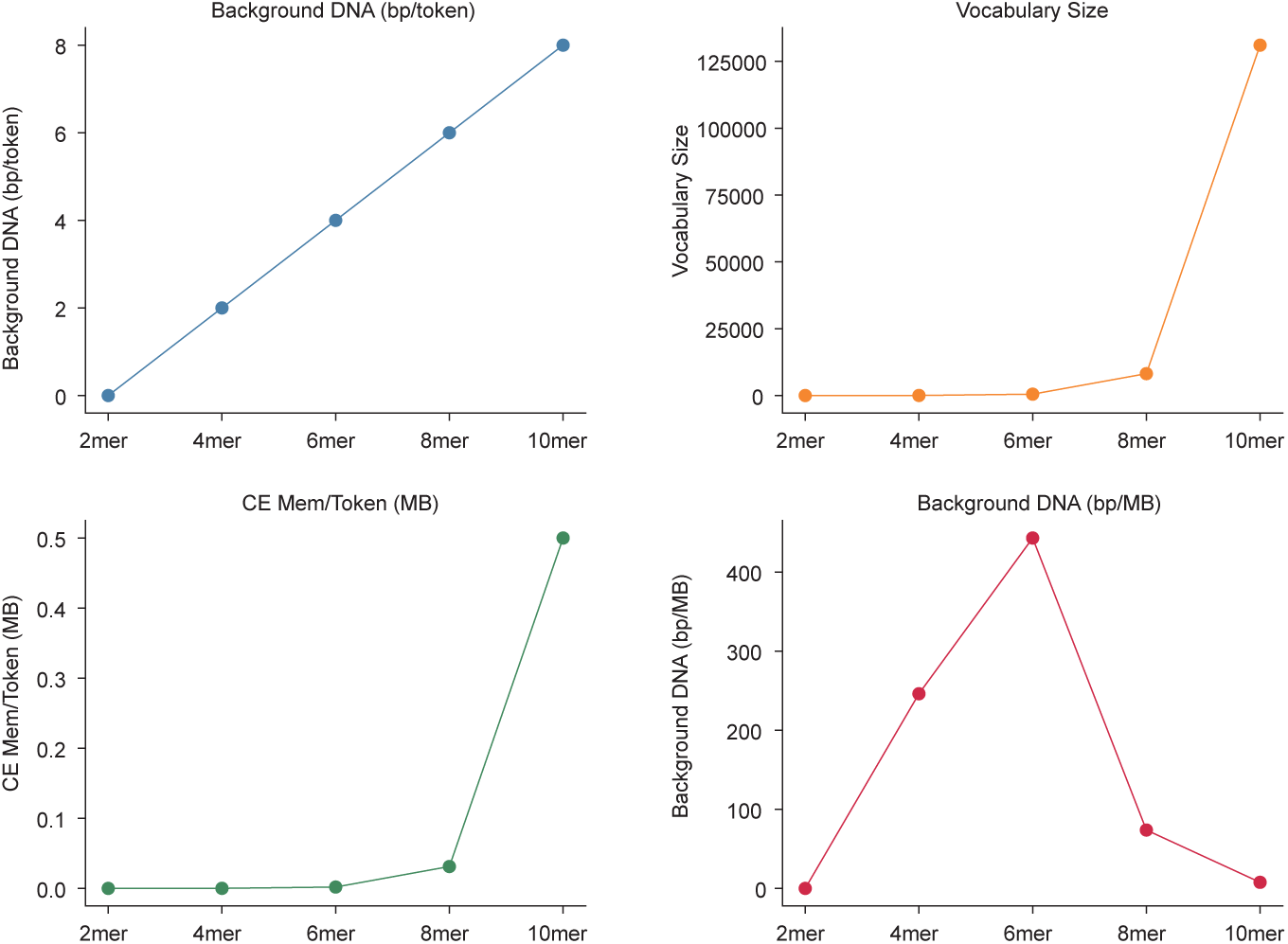
tokenOptimization of token design for background DNA and memory efficiency. (A) Background DNA (bp/token) increases linearly with k-mer size, highlighting the growing sequence context captured by longer tokens. (B) Vocabulary size grows exponentially as k-mer size increases, leading to a rapid rise in computational complexity. (C) Memory usage per token (CE Mem/Token, MB) sharply increases with k-mer size, particularly after 6-mer, due to the expansion of the vocabulary, significantly impacting available memory for sequence input. (D) Background DNA (bp/MB) reaches its maximum at 6-mer, where the token size provides an optimal balance between background DNA representation and memory usage, supporting the most efficient sequence processing without excessive memory demand.

### 1. Choosing CpG-centered 6-mer as the Optimal Tokenization Strategy

The tokenization strategy is crucial for capturing the maximum background DNA context while ensuring efficient memory usage. As illustrated in Figure S1, the amount of background DNA (bp/token) increases linearly with the k-mer size (Figure S1a), allowing longer tokens to capture more surrounding sequence information. However, this increase in token size leads to an exponential growth in vocabulary size (Figure S1b), which in turn causes a sharp rise in memory usage per token (Cross-Entropy Memory per Token, CE Mem/Token, MB) after the 6-mer threshold (Figure S1c>). This increase in memory demand significantly limits the sequence length that can be processed efficiently. At the same time, the amount of background DNA that can be captured per unit of memory (bp/MB) peaks at 6-mer (Figure S1d), offering the best balance between background DNA coverage and memory efficiency. In conclusion, while larger k-mers allow for more background DNA context, their exponentially increasing memory requirements make 6-mer the most effective tokenization choice for capturing the maximum surrounding DNA information without overwhelming computational resources.

**Table S1.** Benchmark datasets used for model evaluation. Summary of classification datasets spanning diverse biological contexts, including cancer, development, and species differences. Each entry reports the number of annotated cell types, the number of training and testing cells, and whether the dataset was included in the pretraining corpus.

| Datasets | #Classes | #Training Cells | #Testing Cells | #In Pretraining |
| --- | --- | --- | --- | --- |
| human adult brain | 15 | 116,643 | 38,881 | Yes |
| Colorectal Cancer | 5 | 895 | 384 | No |
| Early Embryo Dev.<br>(Human) | 10 | 195 | 85 | No |
| Early Embryo Dev.<br>(Mouse) | 14 | 168 | 72 | No |
| Human Brain Dev. | 9 | 4,096 | 1,759 | No |

**Table S2.**
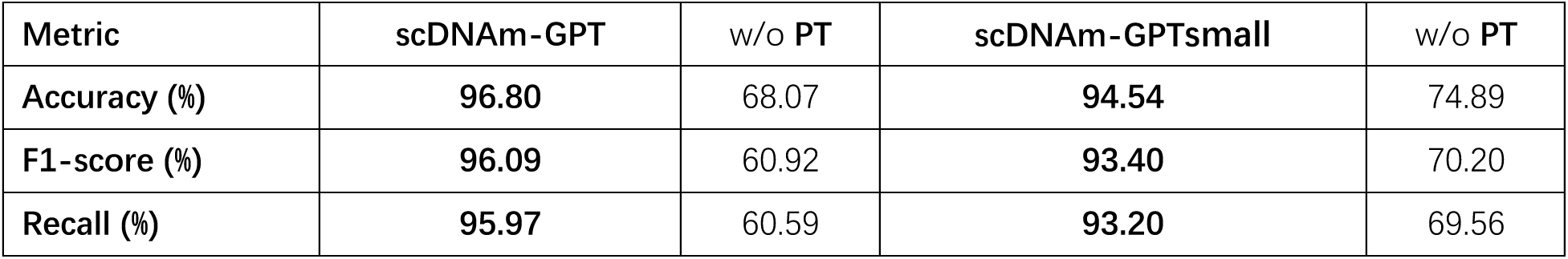
Performance on the human adult brain (seen in the pretraining dataset). scDNAm-GPT is the full pretrained model; w/o PT indicates a randomly initialized version trained from scratch; scDNAm-GPTsmall denotes a 4-layer compact variant. This setting evaluates the model’s memorization and training capacity on data seen during pretraining.

**Table S3.** Performance comparison of scDNAm-GPT and its variants on unseen datasets. We report accuracy, F1-score, and recall across four biological contexts. We compare the scDNAm-GPT (8-layer) and a compact scDNAm-GPTsmall (4-layer) with and without pretraining. These results assess generalization to datasets not seen during pretraining.

| Dataset | scDNAm-GPT | w/o PT | scDNAm-GPTsmall | w/o PT |
| --- | --- | --- | --- | --- |
| **Accuracy** |  |  |  |  |
| Colorectal Cancer | 80.47 | 68.49 | 75.00 | 61.20 |
| Early Embryo Dev. (Human) | 87.06 | 72.94 | 75.29 | 74.12 |
| Human Brain Dev. | 85.15 | 26.31 | 79.19 | 30.96 |
| Early Embryo Dev. (Mouse) | 81.94 | 75.00 | 81.94 | 75.00 |
| **Avg.** | **83.66** | 60.69 | **77.86** | 60.32 |
| - |  |  |  |  |
| **F1-score (%)** |  |  |  |  |
| Colorectal Cancer | 81.86 | 69.56 | 75.66 | 62.81 |
| Early Embryo Dev. (Human) | 91.77 | 70.62 | 75.85 | 76.41 |
| Human Brain Dev. | 84.98 | 23.73 | 79.12 | 28.14 |
| Early Embryo Dev. (Mouse) | 83.77 | 67.35 | 82.11 | 65.74 |
| **Avg.** | **85.59** | 57.82 | **78.19** | 58.28 |
| - |  |  |  |  |
| **Recall (%)** |  |  |  |  |
| Colorectal Cancer | 85.47 | 69.55 | 78.30 | 68.65 |
| Early Embryo Dev. (Human) | 91.70 | 71.69 | 78.81 | 77.07 |
| Human Brain Dev. | 85.04 | 26.30 | 79.09 | 30.91 |
| Early Embryo Dev. (Mouse) | 83.83 | 69.40 | 82.10 | 67.82 |
| **Avg.** | **86.51** | 59.24 | **79.58** | 61.11 |

### 2. Comparison of Full and Compact Models with and without Pretraining

To evaluate the impact of model architecture and pretraining on predictive performance, we systematically compared four variants of scDNAm-GPT: the full model, a compact 4-layer version (scDNAm-GPT_small_), and their respective counterparts trained from scratch without pretraining. Performance was assessed on a diverse panel of datasets spanning different biological contexts, including cancer, development, and cross-species generalization (Table S1).

As shown in Table S3, pretrained models consistently outperformed their non-pretrained counterparts across all four unseen datasets and three key evaluation metrics—accuracy, F1-score, and recall—underscoring the critical role of large-scale methylation pretraining. In particular, scDNAm-GPT achieved an average accuracy of 83.66% on the unseen datasets, compared to only 60.69% for the same architecture without pretraining. Similarly, the compact scDNAm-GPTsmall reached 77.86%accuracy with pretraining, far exceeding its randomly initialized counterpart (60.32%).

Importantly, scaling up model depth from 4 layers to the full scDNAm-GPT architecture led to notable performance gains, especially on complex datasets such as Human Brain Development and Colorectal Cancer. On the brain development dataset, for example, scDNAm-GPT achieved 85.15% accuracy, outperforming scDNAm-GPT_small_ by over 6 percentage points, and the performance gap widened dramatically in the absence of pretraining (26.31% vs. 30.96%). These results demonstrate that pretraining and model capacity act synergistically to enable accurate and robust representation learning, particularly under the challenge of sparse single-cell methylation input.

To assess whether the model can retain information from the pretraining distribution, we further evaluated performance on the Human Adult Brain dataset, which was included in the pretraining corpus (Table S2). As expected, all models showed improved performance on this seen dataset. scDNAm-GPT achieved 96.8% accuracy, indicating strong memorization and representation retention capabilities, while its non-pretrained variant reached only 68.07%, suggesting that pretraining not only aids generalization but also preserves high-fidelity encoding of training distributions.

Together, these findings highlight two key conclusions: i) pretraining is essential for learning biologically meaningful patterns from sparse, high-dimensional methylation data; and ii) increasing model depth substantially enhances predictive performance, especially in biologically heterogeneous or epigenetically complex settings.

**Figure S2.**
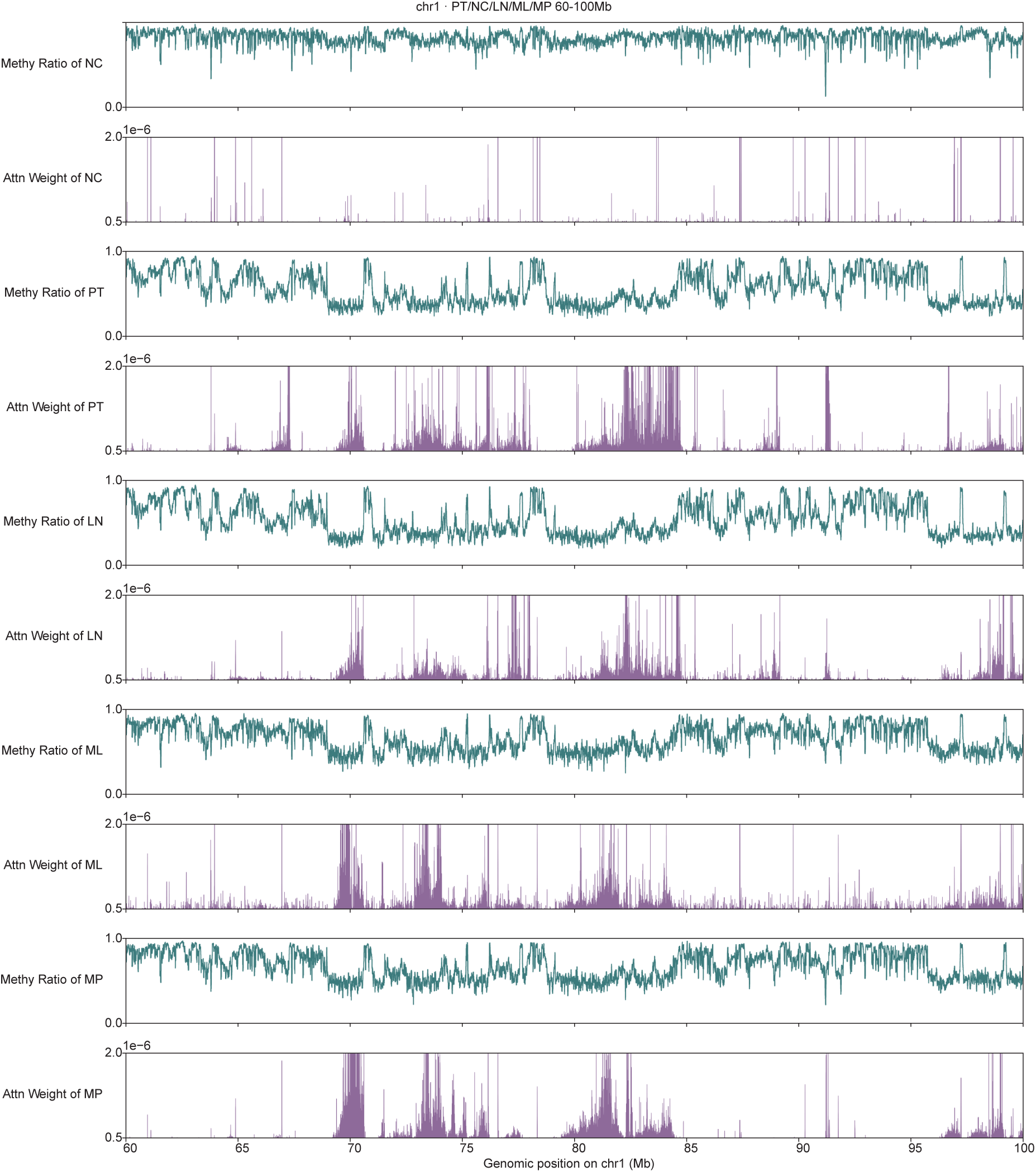
Attention weight patterns of the model across normal colon and various lesions. The model’s attention on Normal Colon primarily exhibits a random, momentary focus. In contrast, for disease tissues, the model concentrates its attention on large regions of hypomethylation, indicating a more focused and consistent pattern of interest.

**Figure S3.**
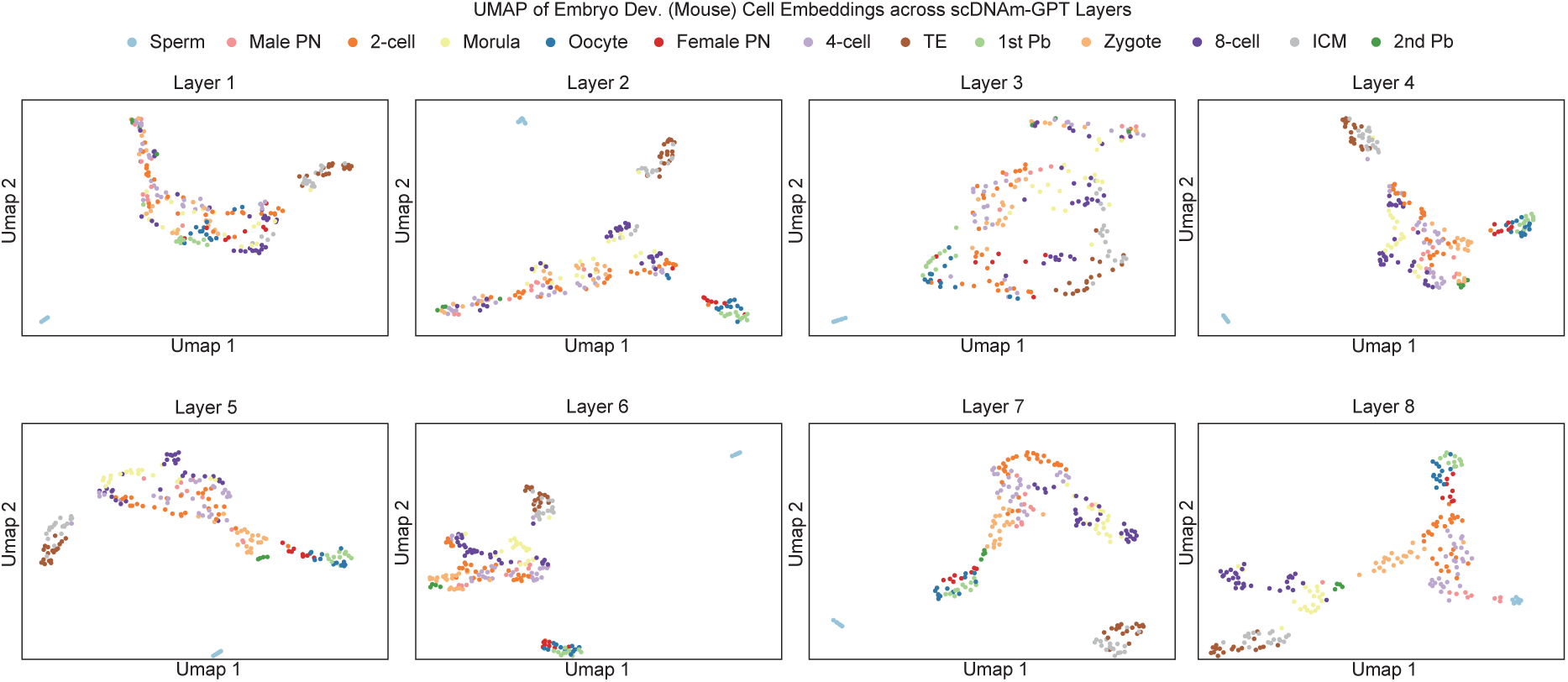
Layer-wise representations reveal hierarchical encoding of developmental signals. UMAP visualizations of scDNAm-GPT embeddings from different layers for mouse early embryo development. Different layers capture distinct types of information, with some layers emphasizing cell-type separation (classification) and others reflecting developmental continuity (pseudotime).

**Figure S4.**
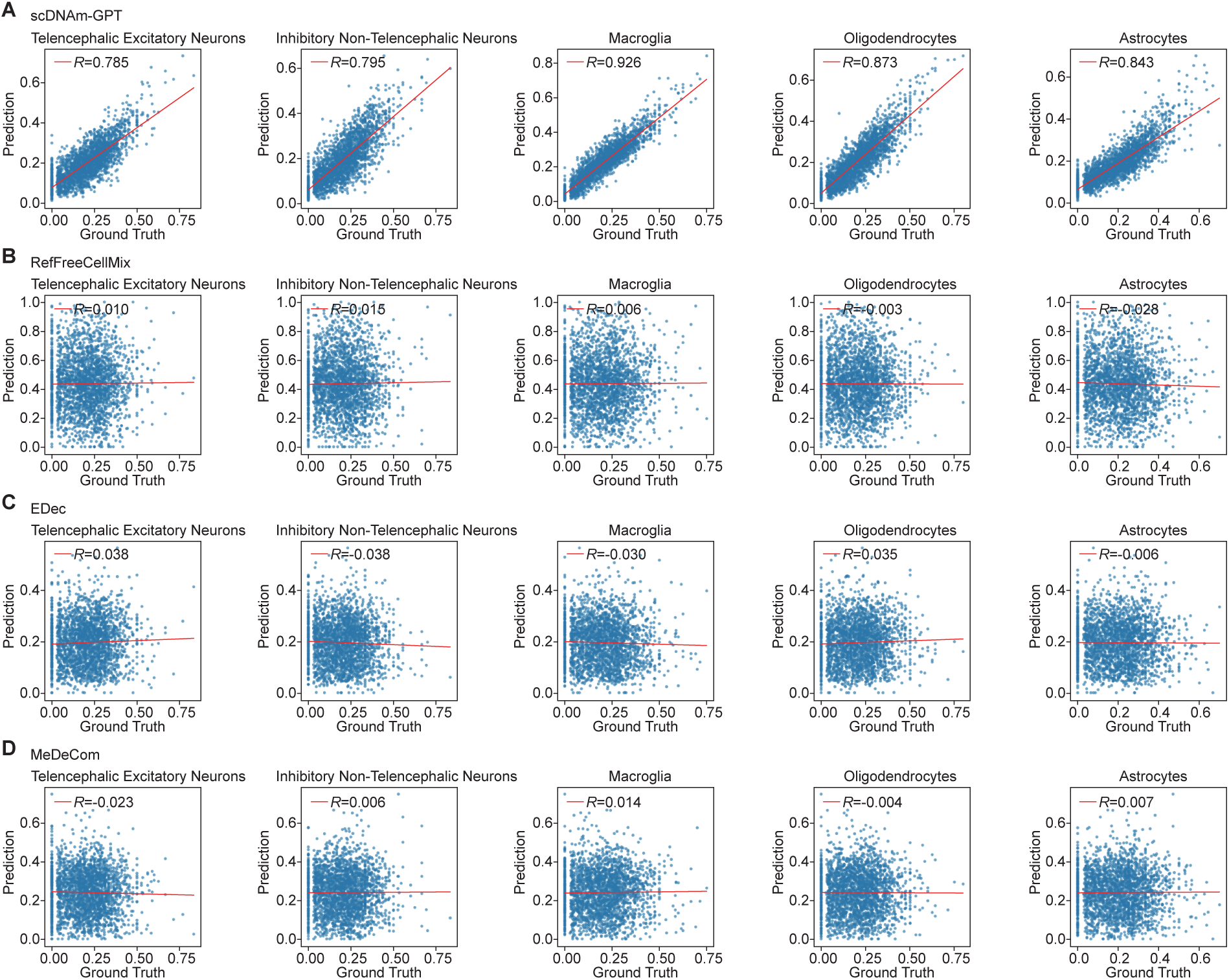
Comparison of reference-free deconvolution methods on neuronal subtype composition inference. (A) scDNAm-GPT achieves an average r = 0.84 (see Figure 6d). (B–D) Estimated cell type proportions from RefFreeCellMix, EDec and MeDeCom show poor concordance with true proportions across five neuronal subtypes (average Pearson −0.05 < r < 0.05). These methods fail to resolve fine-grained methylation differences among closely related neuronal subtypes.

